# Predicting targeted- and immunotherapeutic response outcomes in melanoma with single-cell Raman spectroscopy and AI

**DOI:** 10.1101/2025.05.16.654612

**Authors:** Kai Chang, Mamatha Serasanambati, Baba Ogunlade, Hsiu-Ju Hsu, James Agolia, Ariel Stiber, Jeffrey Gu, Jay Chadokiya, Grayson E. Rodriguez, Prabhjeet Singh, Saurabh Sharma, Amanda Gonçalves, Ojasvi Verma, Fareeha Safir, Nhat Vu, K. Christopher Garcia, Daniel Delitto, Amanda Kirane, Jennifer A. Dionne

**Author notes:** denotes equal contribution (Amanda Kirane, Jennifer A. Dionne). **Prior Presentation.** Presented in part at the 2025 American Association for Cancer Research Annual Meeting, Chicago, IL, April 25-30, 2025.

## Abstract

**PURPOSE:** Identifying reliable predictors of immunotherapeutic response in melanoma remains an outstanding challenge. Existing transcriptomic and proteomic profiling methods for the tumor-immune microenvironment (TIME) are costly and may not faithfully capture modifications actively impacting tumor behavior. Here, we present a non-destructive, single-cell approach combining Raman spectroscopy and machine learning (ML) that enables rapid cell profiling and therapeutic response prediction.

**METHODS:** We analyzed single-cell Raman spectra of mouse and human melanoma cell lines alongside nine melanoma patient-derived samples with known resistance profiles to targeted and immunotherapeutic inhibitors bemcentinib, cabozantinib, dabrafenib, nivolumab, and a combination of nivolumab and relatlimab. We assessed cell phenotyping classification and treatment resistance using random forests and feature importance analysis. For patient samples, we constructed a two-stage evaluation workflow to determine clinical drug resistance through aggregated single-cell predictions and identified corresponding highly variant spectral signatures using computational methods adapted from single-cell RNA sequencing methods.

**RESULTS:** In cell lines, our approach achieved >96% differentiation accuracy across tumor microenvironment cell types and induced functional phenotypes. Persistent (drug-resistant) cells formed subclusters based on genetic mutations rather than sample origin, with Raman signatures reflecting biochemical changes relevant to therapeutic pathways. For patient samples, our workflow correctly inferred resistance likelihoods for 30 of 33 clinically-relevant patient-drug combinations (91% accuracy).

**CONCLUSION:** Single-cell Raman spectroscopy combined with machine learning offers a scalable, prognostic platform to predict therapeutic resistance likelihood, with further potential to advance clinical, multi-omic biomarker efforts for melanoma. Our approach may improve first-and second-line therapy selection assessments for precision medicine by providing rapid, non-destructive prediction of therapeutic response based on cellular spectral profiles.

**Context Summary:** *Key objective:* Can label-free, single-cell Raman spectroscopy and machine learning approach accurately profile melanoma cell states and therapeutic resistance likelihood to targeted and immunotherapeutic agents?

*Knowledge generated:* Raman spectroscopy with machine learning differentiated tumor microenvironment cell types and functional phenotypes with >96% accuracy in cell lines. When applied to patient-derived metastatic melanoma samples, the approach correctly inferred patient response to a panel of targeted and immunotherapeutic inhibitors with 91% accuracy (30 of 33 cases). High-likelihood persistent and sensitive cells across diverse patients exhibited recurrent spectral features.

*Relevance:* Single-cell Raman-based profiling supports functional-diagnostic assessment or resistance likelihood and may contribute to improved therapeutic selection and precision oncology strategies for melanoma patients.

## Introduction

Precision therapies using targeted inhibitors and immunotherapeutic agents have improved outcomes for patients with malignant and metastatic neoplasms. Immunotherapies are the standard first-line treatments for metastatic melanoma, yet demonstrate response rates of <40-50% with a 30-50% risk of severe adverse events.^1–3^ Accurate methods to predict therapeutic response remain a critical unmet need for guiding treatment selection and improving melanoma patient outcomes.^4–7^

Current single-cell transcriptomic and proteomic methods may not always capture structural or post-translational modifications of biomarkers actively impacting response.^4,8,9^ These approaches often require destructive sample processing including cell lysis or antibody labeling, are resource-intensive, and are not readily adaptable to rapid functional assessment. High-throughput, high-resolution, and non-destructive methods capable of functionally profiling tumor cells and forecasting therapeutic response remain highly desirable.

Raman spectroscopy represents an attractive, non-destructive and label-free alternative. By capturing inelastically scattered photons from the molecular vibrations of a sample, spontaneous Raman provides information-dense cell profiles comprising nucleic acids, proteins, metabolites, and lipids. Recently, Raman has demonstrated utility for single-cell phenotyping, from organ-level cell classification to identifying breast cancer subtypes.^10,11^ This technique has also been extended for screening antibiotic susceptibility in tuberculosis and assessing vemurafenib resistance in melanoma cell lines.^12,13^ Continued development of Raman as a complementary functional profiling approach for monitoring tumor and tumor microenvironment responses to therapy represents a major opportunity.

In this work, we profile single cells using Raman spectroscopy and machine learning (ML) to 1) discriminate cell phenotypes commonly found in the TIME, 2) track biochemical changes in murine and human cell lines associated with environmental responses, and 3) determine resistance to targeted therapy and immunotherapy in clinically-relevant melanoma samples. We evaluate the potential of Raman spectroscopy as a functional diagnostic platform to inform therapeutic response prediction in clinical oncology.

## Methods

### Melanoma cell lines

Human melanoma (A375, SK-MEL-24), mice melanoma (YUMM1.7, YUMMER1.7, B16-F10) and mice macrophage cell lines (RAW264.7) were acquired from ATCC (Manassas, VA); SK-MEL-30 human cell line from DSMZ-German collection (Braunschweig, Germany); and LN6-Engleman mouse cell line from the Engleman Lab (Stanford, CA). Cells were cultured in RPMI media with 10% FBS. RAW264.7 cells were stimulated with 100 ng/mL LPS (M1-like) and 20 ng/mL IL-4 + IL-13 (M2-like). Plasticity changes were evaluated by FACS and brightfield imaging. Cell lines were treated with bemcentinib, cabozantinib, dabrafenib, and nivolumab for 24 hours. See Data Supplement for details.

### Human melanoma patient-derived cells

Melanoma tissue specimens were procured from patients undergoing treatment, following the approved tissue collection protocol (IRB 65607) established by the Stanford University Institutional Review Board, with written informed consent. Samples were collected from adult individuals of both genders. All surgical specimens were confirmed to be melanoma prior to processing. Melanoma tissues were collected in RPMI, transported, rinsed with DPBS, and removed of excess connective and vascular tissues. Minced tissues were centrifuged, resuspended, and seeded. Upon confluency, cells were treated with bemcentinib (8 µM), cabozantinib (9.5 µM), dabrafenib (5 µM), nivolumab (6 µM), and nivolumab + relatlimab (6 + 7 µM). scRNA-seq libraries were prepared using 10x Genomics Chromium GEM-X. Libraries were converted and sequenced on Singular Genomics G4.

### Cell fixation for Raman spectroscopy

Cells were isolated, trypsinized, resuspended, and fixed with paraformaldehyde, and monolayers of fixed cells were dropcast on gold-coated glass slides and air dried (Figure 1a). Formaldehyde fixation ensured preservation of cell content for cross-sample spectral analysis.^14,15^

**Figure 1.**
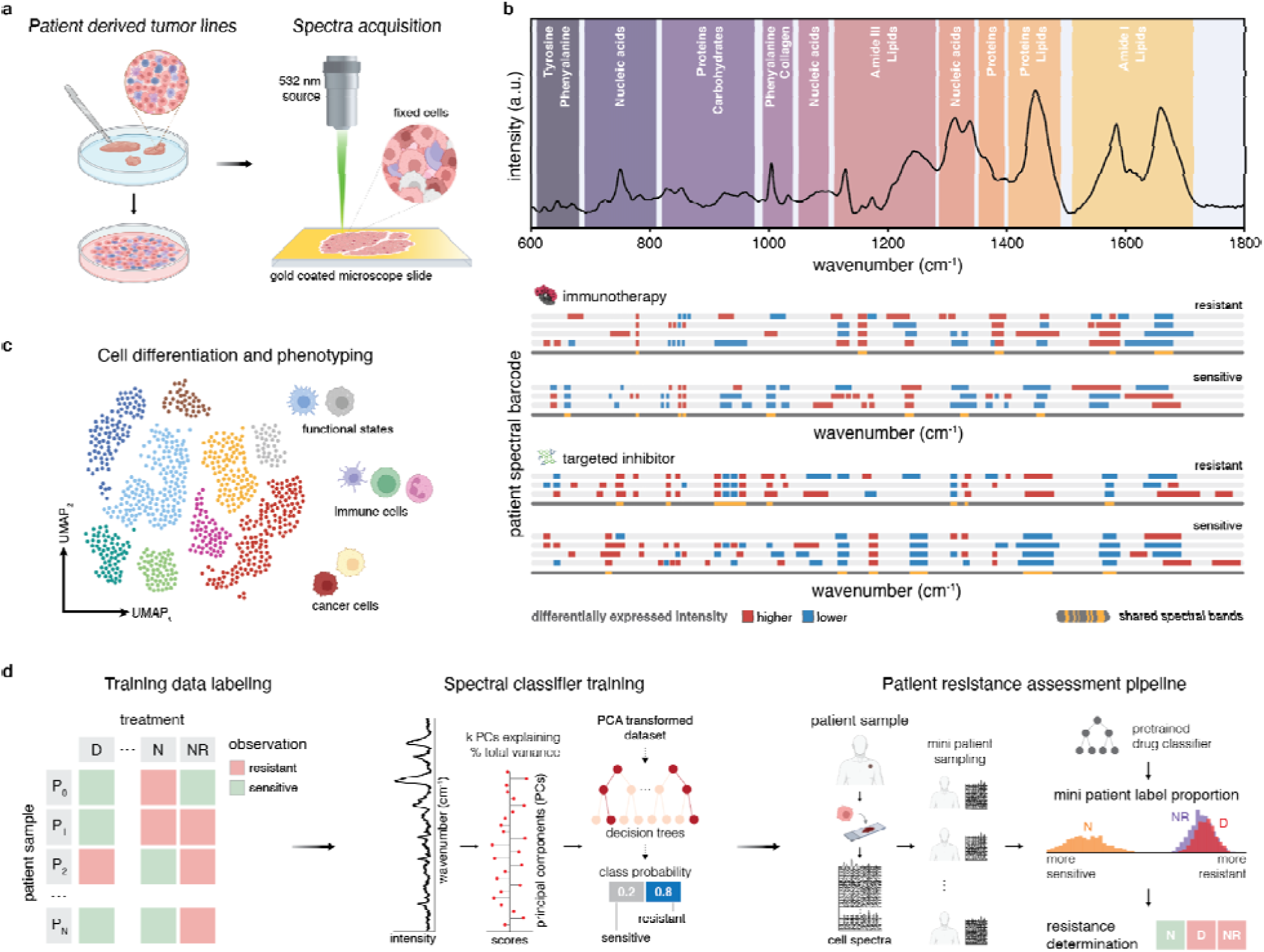
Overview of Raman-based profiling architecture. A) Spectral acquisition overview. Cells from commercial cell lines and patient-derived cell lines were cultured and fixed prior to drop casting onto a gold-coated microscope slide. Spectra are acquired by confocal Raman imaging at a 532 nm excitation wavelength at few– to single-cell resolution. B) Biological Raman window, also known as the optical fingerprint regime, of a cell. Averaged spectra of RAW264.7 macrophages, with major band assignments highlighting cellular makeup. Differential Raman “expression” profiles or spectral barcodes generated from cell spectra are used to delineate samples and drug response. C) UMAP embeddings of melanoma spectra are analyzed by nonlinear dimensionality reduction to examine cell states and phenotypes. D) Patient-derived cell lines resistance determination workflow. Patient clinical response and in-vitro treatment response outcomes are used to label spectral data. Patient-derived cell spectra are collected and reduced using PCA, and a random forest is trained to determine resistance likelihood per spectrum. For an unseen patient, groupings of finite sets of spectra (mini patients) are generated by sampling the patient spectral set. Mini patient label distributions, determined by our pre-trained single spectrum classifier, are used to determine patient resistance to targeted inhibitors and immunotherapies.

### Raman spectral measurements

Spectral acquisition and processing are detailed in the experimental section of the Data Supplement. Briefly, cell spectra were acquired (1000+ per sample) at 532 nm excitation using LabRAM HR Evolution (Horiba). Spectral acquisition approaches and resolution considerations are detailed in the Data Supplement. Preprocessing steps (Supplementary Figure 1) were performed prior to analysis, with spectra isolated at 600-1800 cm^-1^ (Figure 1b). Reference datasets were constructed from >47,000 cancer and immune cell lines spectra and augmented with >27,000 clinical melanoma patient sample spectra.

### Data Analysis

#### Spectral clustering and barcodes

Preprocessed spectra were dimensionally reduced by PCA (capturing ≥95% of variance) prior to UMAP analysis (Figure 1c). Spectral differences among clusters were examined by Leiden clustering (Scanpy^16^). Differential wavenumber “expression” heatmaps were generated from the median log2 fold change with color scale bounds from –2 to +2. Highly variant wavenumbers were identified by (1) selecting the highest feature importance via perturbation analysis (see feature importance) and (2) selecting prominent peaks from the average spectra and from Wilcoxon rank-sum test. Spectral barcodes were constructed from averaged differential intensities to identify biochemical bands indicative of treatment response (Figure 1b). See Data Supplement for details.

#### Spectral classification and feature importance

PCA-transformed spectra were classified by random decision forests (RF, sklearn^17^) with stratified split k-fold cross-validation and uneven class weights. Feature importance was determined by analyzing the classification accuracy dropoff from spectra masking^18^ averaging accuracy as each wavenumber feature is perturbed with a Voigt profile of the pre-transformed data for each test split.

#### Resistance determination architecture

Patient-resistance training matrices were constructed using human cell lines and patients with known clinical (outcome) or *in vitro* therapeutic response (cell viability) (Figure 1d, Supplementary Table 1). For each drug of interest, an RF was first trained on PCA-reduced spectra for single-spectrum resistance prognosis. To determine patient response, single spectrum probabilities were aggregated into subsets known as mini patients. Resistance likelihood was determined from the probability distribution across mini patients. This approach was designed to address intratumoral heterogeneity of metastatic melanoma.

#### scRNA-seq analysis

Alignment was performed with Cell Ranger, and downstream analysis was performed using scanpy and scVI.^16,19,20^ Cells were manually annotated as tumor cells and fibroblast-like cells using canonical marker genes. See Data Supplement for more details.

## Results

### Tumor immune microenvironment cell differentiation

We first assessed cell type and phenotype differentiability using murine melanoma and immune cell lines representing TIME-relevant conditions: distinct cell types (macrophage RAW264.7 and cancer cells), low and high somatic mutations (melanoma YUMM1.7 and YUMMER1.7), and intertumoral heterogeneity (melanoma B16-F10 and LN6-Engleman).^21–23^

We collected an average (μ_cells_) of 1000+ few-to-single-cell spectra per cell line. SEM micrographs confirmed relative monocell density and uniformity (Figure 2a). Mean Raman spectra (Figure 2b) showed distinguishable vibrational peaks (complete assignments in Supplementary Table 4) corresponding to phenylalanine (1003, 1606), nucleic acids (667, 748, 780, 1306, 1337, 1585), protein (748, 1126, 1336, 1446, 1585, 1664), and lipid (1306, 1446 cm^-1^) bands^10,24–27^. Low-dimensional embedding (UMAP) on the top 50 principal components showed YUMM1.7 and B16-F10 groupings (Figure 2b) attributed to the higher melanin presence, which was visible in optical micrographs (Supplementary Figure 2) and in Raman (1385, 1580 cm^-1^).^28,29^ Measurements repeated across multiple days exhibited minimal spectral variability (Supplementary Figure 3). RF achieved 96% classification accuracy discriminating cell lines, with >91% accuracy for parent and derivative lines (Figure 2b).

**Figure 2.**
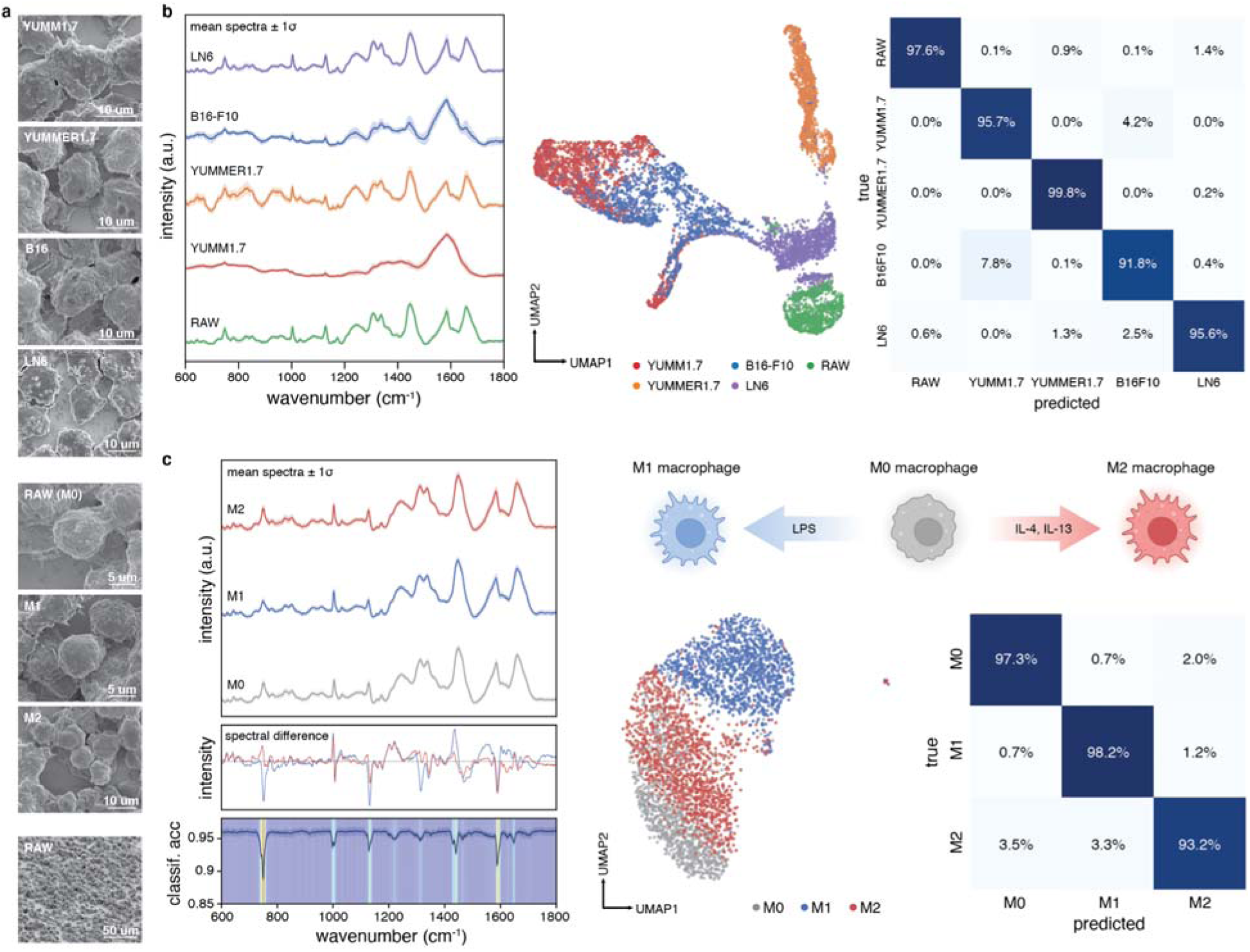
TIME cell type and macrophage polarization state differentiation. A) SEM micrographs of mouse melanoma (YUMM1.7, YUMMER1.7, B16-F10, LN6-Engleman) and macrophage (RAW264.7) cells highlighting minimal morphology differences. The tilted SEM micrograph (bottom) shows cellular aggregation after drying. B) Left: Mean normalized Raman spectra in the biological fingerprint regime of different melanoma and immune cells with ±1 standard deviation. Over 1000 spectra were collected for each cell type. Center: UMAP embedding of PCA-reduced Raman signal. Right: Confusion matrix generated via random forest classification, showing an average of ≥95% accuracy across cell types. C) Left: Mean normalized Raman spectra of RAW264.7 macrophages in different polarization states (M0, M1-like, M2-like) with ±1 standard deviation, spectral differences of polarization states relative to M0 (grey), and wavenumber importance as a function of dropoff in classification accuracy across states. Center: UMAP embedding of macrophage profiles. Right: Confusion matrix showing an average of ≥94% accuracy across polarizations.

### Macrophage phenotype polarization

Having differentiated TIME cell types, we next assessed whether Raman could characterize cell activation and functional phenotype changes relevant in tumor-immune and exogenous agent responses. Inert RAW264.7 macrophages (M0) were exposed to elevated concentrations of cytokines (IL-4 and IL-13) and lipopolysaccharides (LPS) for 24 hours (Figure 2c). Brightfield imaging revealed distinct morphologies: inert M0 (circular), tumor-suppressing M1-like (spindle-like, elongated), and tumor and angiogenesis-assisting M2-like (mixed) plasticities (Supplementary Figure 2), concurrent with prior findings.^30^

Although physical differences (SEM imaging) diminished after sample preparation (Figure 2a), Raman spectra (μ_cells_ = 1198) revealed differences across nucleic acid (749, 782, 1337), lipid (1310, 1655), and protein bands (749, 1003, 1127, 1207, 1447, 1585, 1660 cm^-1^) (Figure 2c).^31–34^ UMAP analysis distinguished M1-like from M0 and M2-like macrophages, with >96% RF classification accuracy. Wavenumber importance analysis identified 750 cm^-1^ (tryptophan) as the primary discriminator (>7% dropoff), followed by protein and lipid bands at 1588, 1440, 1129, and 998 cm^-1^ (Figure 2c), suggesting increased tryptophan metabolic activity positively correlated with M1-like polarization.^35^ These findings establish that Raman can discern environmentally-induced plasticities relevant to therapeutic response.^36,37^

### Melanoma response to therapies in murine and human samples

To determine Raman sensitivity to biomolecular changes impacting therapeutic resistance, we focused on the initial cellular changes in persister cells following short-term drug exposure, which prior work has shown as critical and indicative of long-term resistance development.^38,39^ We investigated murine and human cells pre– and post-treatment with targeted and immune checkpoint inhibitors disrupting critical metastatic melanoma progression pathways. Cells were exposed to IC_50_ concentrations of bemcentinib, cabozantinib, dabrafenib, and nivolumab for 24 hours without medium replacement (Figure 3a), and surviving cells were isolated for measurement. The 24-hour window was selected to minimize confounding factors (cell death, proliferation) while capturing potential early biochemical changes preceding long-term adaptations and resistance.^38^

**Figure 3.**
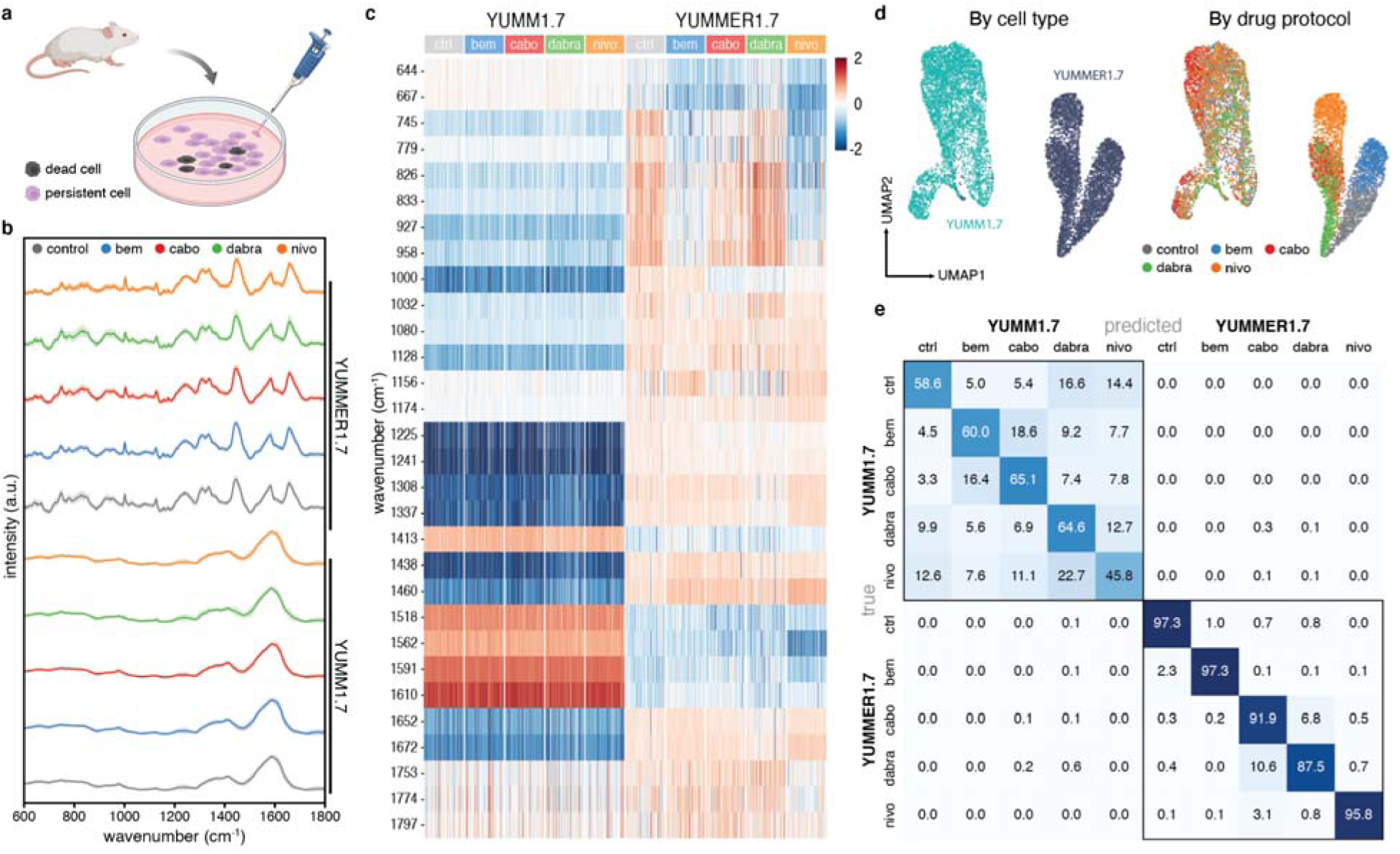
Targeted inhibitor and immunotherapeutic response analysis in murine melanoma cells. A) Drug treatment protocol for commercial murine melanoma cell lines. Cell viability was evaluated for each drug-treated sample, and persistent cells were isolated for Raman spectroscopy. B) Mean normalized Raman spectra and corresponding ±1 standard deviation across YUMM1.7 and YUMMER1.7 cells after exposure to various therapeutic molecules. Over 900 spectra were collected for each sample on average. C) Heatmap of major differential wavenumber bands crucial for cell discrimination, quantified by the median log2 fold change. Wavenumbers are selected from those exhibiting the most significant effect on classification accuracy. Differential Raman intensities are initially stratified by cell type of origin and further by drug protocol. D) UMAP embedding of PCA-reduced spectra from YUMM1.7 and YUMMER1.7 cell response. Left: UMAP grouped by cell type labels. Right: UMAP grouped by drug protocol (color coded grey for untreated, blue for bemcentinib (bem), red for cabozantinib (cabo), green for dabrafenib (dabra), and orange for nivolumab (nivo). E) Confusion matrix for cell type and drug treatment demonstrating higher and lower spectral discernibility for YUMMER1.7 and YUMM1.7, respectively.

In our mouse model, YUMM1.7 and YUMMER1.7 spectra (μ_cells_ = 994, min: 838, max: 1146) divergence and UMAP embeddings agreed with expected responses (Figures 3b,d). Post-treatment YUMMER1.7 displayed more pronounced spectral differences among treatments, likely due to elevated UVB-driven somatic mutations, whereas wild-type YUMM1.7 showed fewer differences^21,22,40^, possibly obscured by prominent melanin-related features (Supplementary Figure 2).

Adapting scRNAseq analysis^41^, we quantitatively assessed cell type-drug spectral disparities by differential Raman “expression” (intensity) analysis (Figure 3c). YUMM1.7 showed minimal changes, whereas YUMMER1.7 exhibited drug-specific changes. Post-bemcentinib YUMMER1.7 cells showed intensity changes at 644, 667-833, 1438, 1460, and 1610 cm^-1^ coinciding with tyrosine assignments^42–44^, and elevated intensities at carotenoid and protein bands (1156 cm^-1^). Post-cabozantinib showed similar changes plus a decrease at 1000 cm^-1^. Flow cytometry and BCA protein quantification on independent samples substantiated the treatment-induced biochemical changes seen in Raman. Post-bemcentinib cells had significantly elevated neutral lipids (p<0.05) with corresponding key lipid-associated spectral changes, while both treatment groups showed decreased protein content (p<0.05) concordant with intensity decreases at protein-specific bands (Supplementary Figure 4)^45–52^. Statistical analysis using Tukey’s HSD and linear mixed-effects models with FDR correction identified treatment-specific molecular signatures concordant with orthogonal biomolecular data. Temporal analysis of post-bemcentinib cells at 30 minutes and 24 hours revealed pronounced early changes across multiple spectral bands with partial recovery toward baseline by 24 hours. This trajectory is consistent with acute response followed by adaptive remodeling in drug-tolerant states, with adaptation initiating within 24 hours of exposure (Supplementary Figure 5).^53–57^

Dabrafenib-persistent YUMMER1.7 showed changes at 745-958, 1000, 1032, and 1337 cm^-1^, in alignment with their BRAF inhibitor sensitivity.^58,59^ No observed major peaks corresponded to relevant inhibitor spectra (Supplementary Figure 6). Nivolumab-persistent YUMMER1.7 showed marked changes in protein (745, 826, 927, 1460, 1562, and 1672 cm^-1^) and nucleic acid (745, 779, and 833 cm^-1^) bands, which may reflect the presence of PD-1 variants (iPD-1) unique to YUMMER, resulting in cell cytostasis.^60,61^ Our classifier achieved >99.1% isolate-level accuracy across and 76.4% within cell types (Figure 3e). Detailed cell-drug one-versus-all groupings can be found in Supplementary Figure 7.

Next, we investigated Raman spectra (μ_cells_ = 1357, min: 336, max: 3327) from human commercial melanoma cell lines (A375, SK-MEL-24, and SK-MEL-30) and patient-derived tumor lines (PAT-52 and PAT-73). While untreated samples showed broadly different spectral signatures, similarly treated persistent cells displayed intensity change affinities and formed subclusters based on relevant genetic mutations rather than sample origin (Figure 4a-c, Supplementary Figure 8).

**Figure 4.**
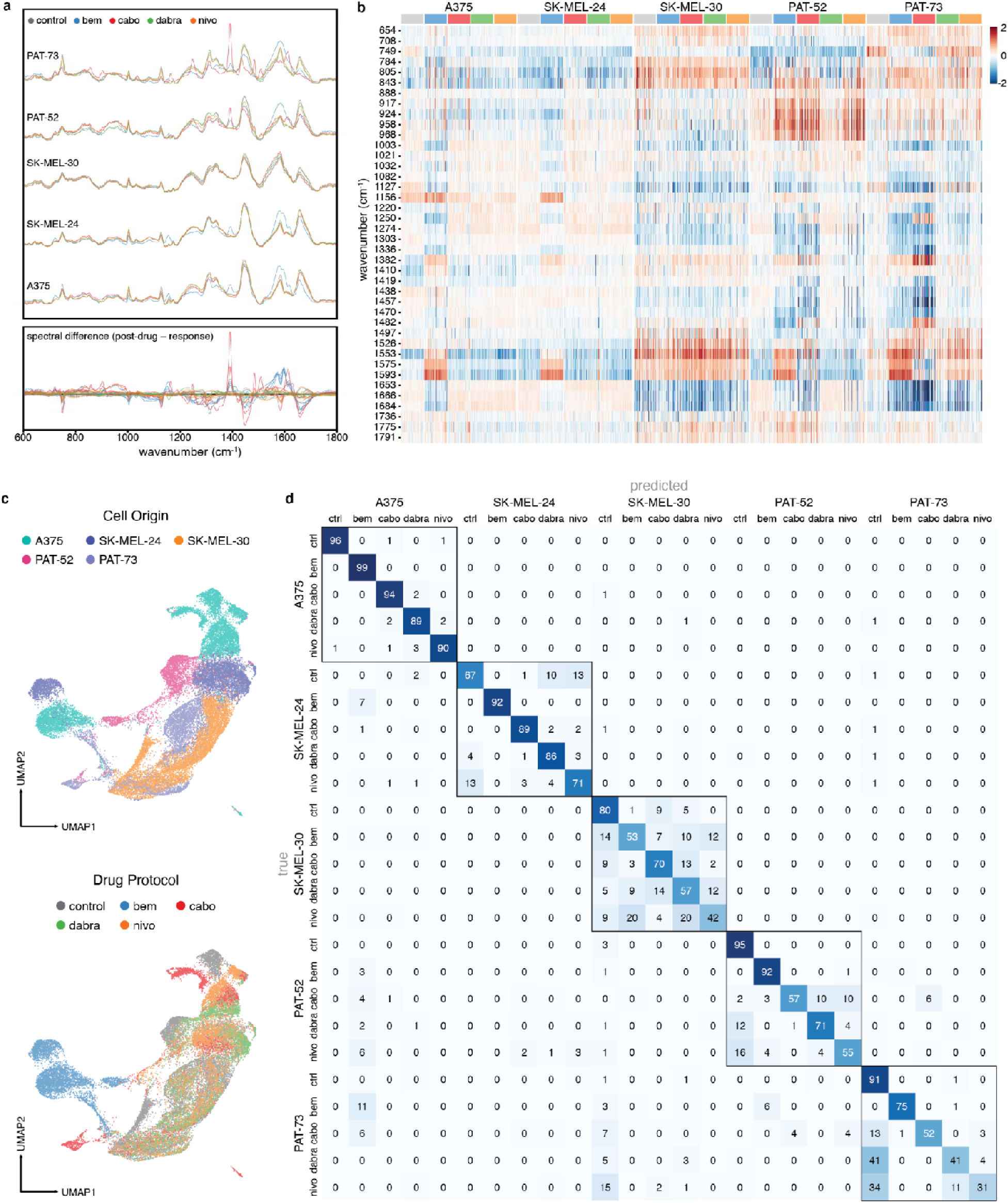
Targeted inhibitor and immunotherapeutic response analysis in human melanoma cell lines and patient-derived cell lines. A) Top: Mean normalized Raman spectra of cell response to treatment protocols, grouped and overlaid by patient-derived cell lines. Bottom: Spectral differences of post-treated sample relative to untreated (control), highlighting the major bands shared by similar treatment protocols across samples. B) Heatmap of differential wavenumber bands expressed in surviving cells, quantified by the median log2 fold change. C) UMAP embedding of Raman profiles from human cell responses. Top and bottom spaces are grouped by cell type labels and drug protocol, respectively. D) Confusion matrix across cell type and drug treatment classification, demonstrating higher inter-sample differentiation and varied treatment discrimination intra-sample.

For example, AXL+ samples (A375, SK-MEL-24) showed post-bemcentinib intensity shifts distinguishable by fold change and unified latent space groupings (Figure 4b). Decreases in protein (1000, 1250, 1336, 1438, 1660 cm^-1^) and lipid (1081, 1438, 1660 cm^-1^) bands likely reflect tyrosine activity and AXL pathway modifications. Meanwhile, persistent AXL– cell lines (SK-MEL-30) were indiscernible from control (Figure 4c). Similarly, post-dabrafenib wild-type BRAF samples (SK-MEL-24, SK-MEL-30) exhibited little change, whereas BRAF+ cells (A375) showed decreases at 749, 1127, and 1526 cm^-1^, in addition to the 1156 and 1582 cm^-1^ changes observed by Qiu et al. for BRAF inhibitor vemurafenib.^13^ Post-cabozantinib cells showed decreased protein and lipid bands at 749 and 1653-1684 cm^-1^ and increases between 800-1000 cm^-1^. Post-treatment PAT-52 and PAT-73 cells saw comparable changes across targeted inhibitors but differed with nivolumab. Post-nivolumab PAT-52 cells had increased protein bands at 843, 917, 924, 958, and 968 cm^-1^, whereas PAT-73 showed minor changes, reflecting its clinical non-responder status.

Comparison of persistent cell spectra with targeted inhibitor spectra suggested remnant inhibitor presence. Increased intensities of post-bemcentinib cells at 1156 and 1550-1600, and a discontinuity at 1615 cm^-1^, coincided with major bemcentinib bands (Supplemental Figure 3). Similarly, a subset of post-cabozantinib cells, isolated by Leiden clustering, exhibited pronounced features at 1388, 1439, 1482, and 1611 cm^-1^, likely corresponding to major cabozantinib peak assignments (Supplementary Figure 8). These findings seemed consistent with cabozantinib (∼120 hours) and bemcentinib (∼150 hours) half-lives.^62,63^

Our classifier retained 94% accuracy preserving patient discernibility (Figure 4d). Within patients, classification generally performed better for those with relevant positively expressed mutations (AXL+, BRAF+ in A375) than for low and wild-type expressions (AXL-, BRAF WT in SK-MEL-30) or nonresponsive (PAT-73) samples. Additionally, bemcentinib and cabozantinib-treated cell spectra appeared similar across patients, respectively and were more likely misclassified. These findings indicate Raman can discern comparative degrees of resistance across diverse cell origins and identify conserved spectral features shared among persistent states. These biochemical signatures provide proof-of-concept towards analyzing clinically-derived samples where intratumoral heterogeneity is expected.

### Patient melanoma sensitivity screening

Finally, we applied our methodology to develop drug determination classifiers for clinically-relevant patient samples. We designed models to identify spectral markers indicative of clinical resistance while accounting for potential cell population variance within patient-derived samples. Building on our previous findings, we investigated whether we could distinguish resistant-associated spectral patterns from patient-specific variations. RF classifiers evaluated resistance-sensitivity profiles for targeted therapies (bemcentinib, cabozantinib, dabrafenib), single-agent (nivolumab) and combination-agent (nivolumab + relatlimab) immunotherapy. We prospectively collected heavily-pretreated resistant patients undergoing surgical resection and pretreatment specimens from patients planned for neoadjuvant therapy, following clinical response. Our dataset encompassed five training and four testing patients (μ_cells_ = 522, min: 96, max: 1512). Additionally, we collected spectra across multiple passages from select patients to examine passage drift. Given limited fresh tissue available prior to therapy, we trained with at least two resistant and sensitive patients per protocol when possible to maximize labeling power while moderately accounting for patient melanoma diversity (Supplementary Table 1).

UMAP visualization highlighted significant melanoma heterogeneity (Figure 5a). Primary tumor-derived cells were mainly consolidated in one region, whereas immunotherapy-refractory (IR) samples showed broader heterogeneity (Figure 5b). Some IR patient spectra (PAT-03, PAT-73) overlapped with commercial cell line spectra, suggesting biological similarities.

**Figure 5.**
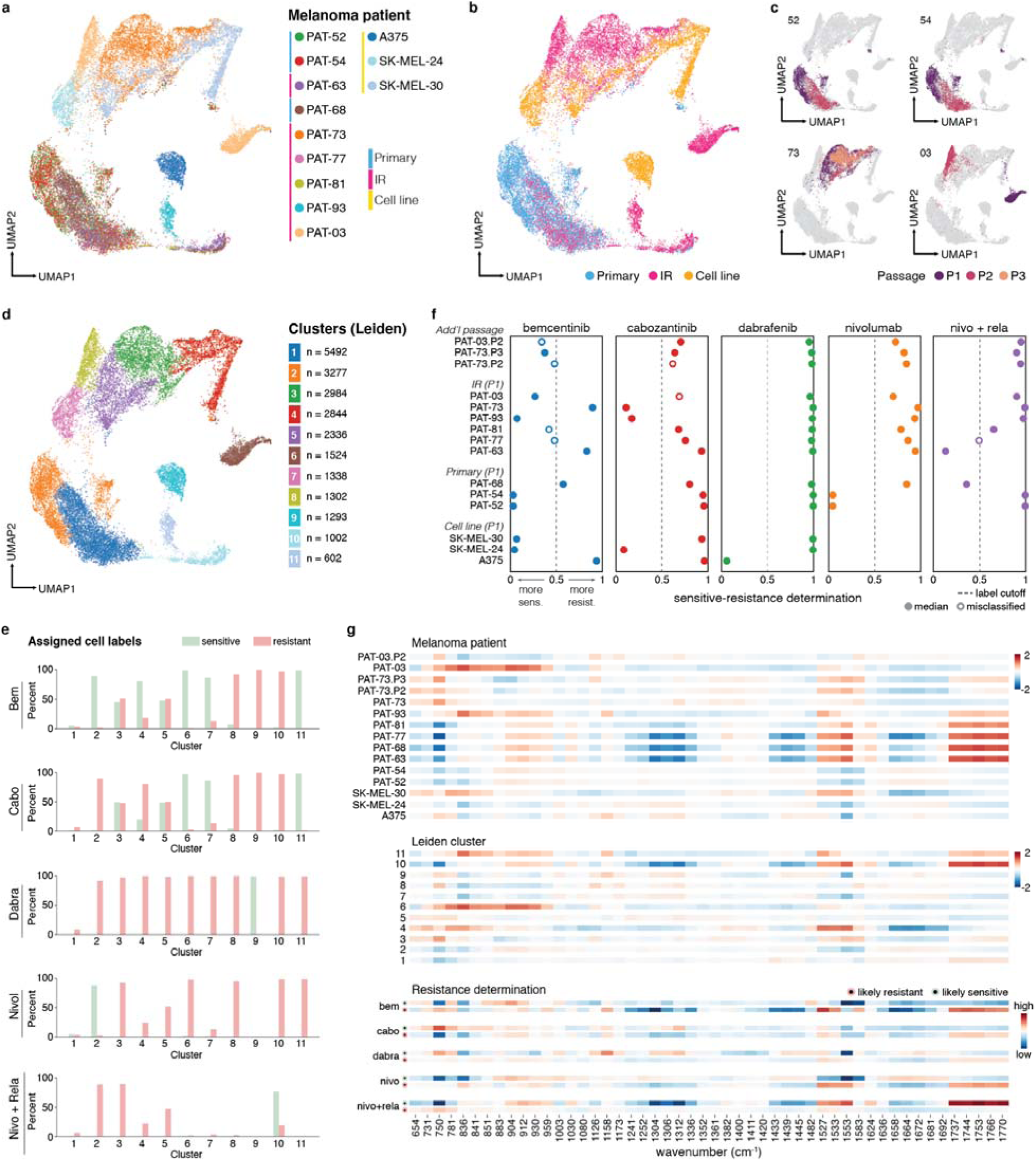
Analysis and response determination of patient-derived cell lines. A) UMAP embedding of Raman profiles from patient-derived cell lines and commercial human cell lines, where each point represents a cell spectrum, showing cellular heterogeneity across patient samples. B) UMAP colored by sample origin. C) UMAP highlighting cell spectra from select patients colored by passage number. D) UMAP embedding colored by Leiden clusters. E) Proportion of cell spectra within each Leiden cluster labeled resistant or sensitive across drugs of interest. F) Response determinations for patient-derived cell lines (seen and unseen by our ML model) across targeted and immunotherapy regimes. Using a 0.5 cutoff boundary to label resistant and sensitive samples, 26 of 33 test drug-patient cases were correctly labeled. G) Spectral barcode of highly variant wavenumbers quantified by the median fold change across samples (top), Leiden clustering (middle), and higher (>0.8) and lower (<0.2) resistance likelihood determined by our ML model (bottom).

We observed tumor compositional drift and cell population convergence across passages, marking phenotypic drift in metastatic progression (Figure 5c). UMAP showed later passages of several patients (PAT-52, PAT-54, PAT-63, and PAT-68) converging to a single grouping different from their initial passages (Figure 5c), with elevated intensities at 1658 and decreased intensities at 750, 1312, and 1583 cm^-1^ (Supplementary Figure 9,10). Meanwhile, later passages of PAT-73 and PAT-03 localized within prior passages or near melanoma cell lines. Single-cell RNA sequencing (scRNA-seq) on later passages revealed near-homogenous fibroblast-like populations for PAT-52, PAT-54, PAT-63, and PAT-68, whereas PAT-73 and PAT-03 showed fewer fibroblast-like genes, consistent with spectral observations (Supplementary Figure 11).

Leiden clustering (resolution=0.5) further identified cell type clusters largely independent of patient origin, in agreement with scRNA-seq (Figure 5d, Supplementary Figure 12). Cluster 1 had a fibroblast-like profile, while clusters 3-5 contained mixtures of patient and commercial tumor cells (Figure 5d). Interestingly, when mapping patient response data (determined by clinical and cell viability assessment) to these clusters (Figure 5e), we found many clusters exhibited homogenous resistance labeling, with similarly grouped patients tending to share drug response profiles. For example, PAT-52 and PAT-54 cells mostly resided in cluster 2 with identical resistance across cabozantinib, dabrafenib, and nivolumab + relatlimab combination (Supplementary Table 1). Similarly, PAT-63 and PAT-68 largely populated cluster 10 and exhibited resistance to all drugs except our combination regimen. These observations suggest Raman profiling may identify similarly responding cells and inform second-line treatment selection.

Using a 0.5 threshold and 25,000 mini patients per patient, our patient resistance likelihood classifier model correctly classified 79% (26/33) of patient-drug combinations (Figure 5f). Our worst performing model (bemcentinib, 3/7 patients) had most misclassified determinations near the cutoff threshold. Adjusting to optimized labeling cutoffs per treatment improved accuracy to 91% (30/33) (Supplementary Figure 13).

Our model also showed passage-associated differences in drug resistance. For example, PAT-03 bemcentinib resistance likelihood increased with passages while PAT-73 resistance likelihood decreased, consistent with cell viability data (Supplementary Table 1). Further analysis showed substantial cluster assignment changes among multiple major subpopulations (Supplementary Figure 14,15).

Spectral barcodes compared across patient samples, Leiden clusters, and highly resistant (likelihood >0.8) or sensitive-labeled (likelihood <0.2) cells revealed treatment-specific differentially expressed bands, notably at 750, 836, 1527, 1553, and 1737-1770 cm^-1^ (Figure 5g). Cross-examination of barcodes provided a more comprehensive response assessment. For example, comparing borderline bemcentinib determinations (PAT-77, PAT-81) with bemcentinib-response barcodes and cluster 10 (predominantly bemcentinib-resistant) adjudicated them as likely resistant. Conversely, PAT-77 remained misclassified for combination immunotherapy, indicating training set size limited improvement. With larger patient sets, this two-stage evaluation approach may better improve second-line therapy selection and reduce toxicity risk from ineffective treatments.^64^

## Discussion

In summary, we present Raman spectroscopy combined with machine learning as a functional, single-cell profiling platform capable of detecting biochemical adaptations associated with therapeutic response. We demonstrate discrimination of TIME cell phenotypes, identification of conserved biochemical features associated with drug-tolerant persister states, and prediction of therapeutic resistance likelihood in clinically relevant melanoma samples. By integrating *in vitro* and patient-derived data, our models correctly infer resistance likelihood for a majority of unseen patient-drug combinations without necessitating post-treatment molecular characterization. These findings support the potential of Raman spectroscopy to functionally profile tumor cells and complement existing molecular approaches. Notably, spectral features associated with resistance were detectable prior to overt phenotypic changes, highlighting the potential of Raman spectroscopy to identify early adaptive states.

Although our results are promising, several limitations warrant consideration. First, the relatively small patient training cohort may have introduced sampling bias and limited generalizability. Expansion of the spectral reference library, particularly with pre-treatment clinical specimens collected through multi-institutional collaborations, would improve model robustness, enable deep learning utilization, and better capture the diversity of melanoma phenotypes. Second, our binary resistance labeling framework simplifies the complex and continuous spectrum of therapeutic response.^65,66^ Future studies incorporating graded response classification, longitudinal sampling, and organoid-based modeling may provide nuanced representations of resistance biology. Third, while fixation improved spectral reproducibility and cross-sample comparability, aldehyde crosslinking introduced expected spectral shifts relative to live cells (Supplementary Figure 16).^67^ Live cell Raman analysis represents an important future direction for capturing dynamic cellular adaptations though technical challenges including background signal variability and laser-induced perturbation must be addressed.

We also note that while Raman spectroscopy provides biochemical insights and our orthogonal validation confirmed treatment-associated lipid and protein alterations, additional targeted molecular investigations may elucidate the specific pathways contributing to these spectral signatures. Finally, although our 24-hour exposure model captures early adaptive changes associated with drug tolerance^39^, extended longitudinal studies would be important for characterizing the full temporal evolution of acquired resistance.

Future technological developments may further enhance the clinical and research utility of this platform. Surface-enhanced Raman spectroscopy (SERS) and nanophotonic substrates may improve sensitivity and throughput^68–71^ i, while spatially resolved Raman profiling offers potential to interrogate intact tumor architecture, tumor-immune interactions, and organoid models.^41^ Integration with advanced machine learning approaches and complementary molecular assays may enable more comprehensive characterization of tumor functional states.

Overall, Raman-based functional profiling represents a rapid, label-free approach for assessing therapeutic resistance likelihood in melanoma. As a non-destructive optical method, Raman spectroscopy offers a complementary modality to existing genomic and proteomic technologies and may contribute to multi-modal precision oncology strategies (e.g. sc-seq, iFRET).^72^ With further validation in larger clinical cohorts, this approach improved therapeutic stratification and response prediction in melanoma and other heterogeneous malignancies. ^72^

## Acknowledgements

The authors are grateful to H. Balch, V. Dolia, H. Carr Delgado, L. Herndon, J. Hu, D. Omo-Lamai, E. Wagner, C. Luna, O. Verma, A. Georgiadis, and Prof. B. Larijani for their insightful discussions. The authors also thank the Reticker-Flynn lab for their assistance with sample procurement. Select elements of Figures 1-3 were created with Biorender.com. J.A.D. is a Chan Zuckerberg Biohub – San Francisco Investigator.

## Contributions

K.C., M.S., A.K., and J.A.D. conceived and designed the experiments. M.S., H.H., S.S., and J.C. contributed to commercial cell and patient sample culturing and preparation. K.C., B.O., and M.S. developed the experiment protocols. K.C., M.S., B.O., and A.S. contributed to sample preparation and performed the spectroscopy measurements. K.C. and A.S. conducted the scanning electron microscopy characterizations. M.S. and H.H. conducted the cell viability measurements. M.S., H.H., S.S., and J.C. carried out the immunofluorescence and optical microscopy characterizations. M.S., H.H., and A.G. contributed to library preparation and performed the single-cell sequencing. K.C. and J.A. analyzed the collected experimental data. N.V., F.S., and J.G. assisted in the development of the feature importance and classification workflow. A.S. assisted in the development of the Raman spectral preprocessing pipeline. J.G. assisted in the development of the outlier removal protocol. G.E.R. performed and analyzed orthogonal intracellular lipid, total protein, and phosphorylation quantification experiments. D.D. assisted in the single-cell sequencing analysis. K.C., M.S., J.A., J.G., D.D., A.K., and J.A.D. contributed to the interpretation of the results. All authors helped shape the research, analysis, and contributed to the manuscript. A.K. and J.A.D. conceived the idea and supervised the project, along with D.D. and K.C.G. on relevant portions of the research.

## Data and code availability

The data that support the findings in the work are available in the article and the supplementary information file, and are available from the corresponding authors on reasonable request. The source code for the analysis and visualization in this study is available from the corresponding authors on reasonable request.

## Notes

**Support.** J.A.D. is a Chan Zuckerberg Biohub – San Francisco Investigator and acknowledges funding from the Chan Zuckerberg Biohub San Francisco, as well as from the National Institutes of Health (NIH) under grant no. 1DPAI15207201. A.K. acknowledges funding from the John and Marva Warnock Endowed Scholar Fund. J.A.D. and A.K. acknowledge funding from the Stanford Cancer Institute (SCI) Innovation Award. B.O. was additionally supported by the Stanford Graduate Fellowship, the Stanford EDGE Fellowship, and the Stanford Bio-X Bowes Graduate Student Fellowship. J.A. acknowledges funding from the Stanford Transplant and Tissue Engineering (TTE) Center of Excellence. A.S. was additionally supported by the National Science Foundation (NSF) through the Graduate Research Fellowships Program. G.E.R. was supported by the Mechanisms and Innovations in Cardiovascular Disease T32 Training Grant (T32EB009035). K.C.G. acknowledges funding from Howard Hughes Medical Institute and the Parker Institute for Cancer Immunotherapy. Part of this work was performed at the Stanford Nano Shared Facilities (SNSF), supported by the National Science Foundation under award ECCS-2026822.

### Competing Interest Statement

The authors have declared no competing interest.

### Summary of Updates

Revisions include additional data and figures in supplemental files reflecting biological validation and methods justification. Main text has additionally been condensed. Authors and funding sources have also been updated. Separated supplementary and main text documents.

