## Supplementary material for "Predicting targeted- and immunotherapeutic response outcomes in melanoma with single-cell Raman spectroscopy and AI"

### Supplementary Methods: Experimental

#### Melanoma cell lines

Cells were cultured in complete cell culture growth medium including RPMI (Gibco, Grand Island, NY) supplemented with 10% fetal bovine serum (Corning, Woodland, CA) at 37°C in an incubator supplied with 5% CO2. Media was changed at least twice a week. Cells were cultured at 37°C in a humidified CO2 incubator at 5% CO2. Cell lines not used in drug response were cultured until 65-70% confluency. Cell lines used in drug response were cultured to 75-80% confluency, trypsinized (Trypsin-EDTA, Gibco) for 5 min at 37°C, collected in suspension in a 15 mL tube, and centrifuged at 400 g for 5 minutes at 4°C. The supernatant was removed, and the cell pellet was resuspended with 1 mL RPMI complete media. Cells were counted and plated at a density of 1x106 cells/well into a 6-well plate and placed overnight in an incubator for adherency. Cells were then treated with desired inhibitors with varying concentrations for 24 hours. Inhibitor concentrations for all samples can be found in Supplementary Table 2. Duplicate wells were used for both treated and untreated cells. Plates were incubated for 24 hours. Dead cells along with the drug media were removed, and cells were trypsinized for 5 minutes, collected into 15 mL tubes, and centrifuged at 400 g for 5 minutes. After supernatant removal, cells were resuspended into 1 mL of media, and cell viability was assessed with Trypan blue assay. Hits were identified using a threshold of 65-70% inhibition of cell viability. For temporal studies, cells from the same batch were split equally and treated with bemcentinib for 30 minutes or 24 hours prior to Raman analysis.

#### Human melanoma patient-derived cells

Melanoma tissues were collected in RPMI media and transported to lab on ice and rinsed with DPBS (Gibco) and removal of excess connective and vascular tissues. Subsequently, tissues were finely minced using a sterile scalpel and collected in the tube. After centrifugation (5 min at 4°C, 400 g) the cell pellet was resuspended in DMEM/F12 (Invitrogen, Waltham, MA), 12 mM HEPES (Invitrogen), 1% GlutaMAX (Invitrogen), 10% fetal bovine serum (FBS), supplemented with 9 µM ROCK (Y-27632) inhibitor (Selleckchem, Houston, TX) and 1% penicillin/streptomycin (Invitrogen). Cells were seeded in a cell culture dish and kept at 37°C and 5% CO2 in an incubator. Once they reached 65-70% confluence, cells were detached with TrypLE Express (Gibco), centrifuged (5 min at 4°C, 400 g) and seeded in 6 well plates at a density of 1x106 cells per well for drug treatment. For select patient samples, persistent cells were further partitioned for fixation protocols used in Raman data acquisition and for preparation for single-cell transcriptomic sequencing.

#### scRNA-seq cell and library preparation

Patient-derived melanoma cell lines were trypsinized for 5 minutes, collected into 15 mL tubes, then spun down at 400 g for 5 min at 4°C. After removing the supernatant, the cells were resuspended in 1 mL of PBS with 0.5% BSA. Live and dead cells were visualized and counted using Trypan blue. If over 40% of dead cells were present, the sample was spun down at 400 g for 5 min at 4°C and subject to the dead cell removal kit (Miltenyi Biotec, 130-090-101). Finally the samples were spun down at 400 g for 5 min at 4°C and resuspended in 500 µL - 1 mL of PBS for a final concentration of 1000-1,500 cells/µL for a target cell recovery of 20,000 cells. For the characterization studies, library preparation was performed using Chromium GEM-X Single Cell 3' Reagent Kits v4 (10x Genomics). Library conversion was then done using Singular Genomics Library Compatibility Kit (Singular Genomics). Libraries were sequenced on the Singular Genomics G4 sequencer, using 100 cycle F3 flow cells (Singular Genomics), with two lanes per library at 28/90 paired end.

**Intracellular neutral lipid staining of live cells**

Approximately $10^5$ YUMMER1.7 cells per sample were harvested and washed with FACS buffer, PBS (20012-050, Gibco) + 0.2% BSA (A9418, Sigma), twice. Cells were stained with 2 µM BODIPY 493/503 (D3922, Thermo) for 30 min at 37°C. Cells were again washed twice with FACS buffer and then resuspended in 1 µg/mL DAPI (D1306, Thermo). Mean fluorescence intensity (MFI) of BODIPY signal was quantified on a CytoFLEX S Flow Cytometer (Beckman Coulter) by gating on live (DAPI-) single cells. Comparison of multiple groups was performed by using one-way analysis of variance (ANOVA) with Dunnett correction.

**Intracellular protein and phosphorylation quantification**

$6x10^6$ YUMMER1.7 cells per sample were lysed on ice for 45 minutes with RIPA Lysis and Extraction Buffer (8990, Thermo) supplemented with Vanadate (P0758L, New England Biolabs), Halt Protease Inhibitor Cocktail (#78430, Thermo), Phosphatase Inhibitor Cocktail I (AB201112, Abcam), and Protease Inhibitor Cocktail (P8849, Sigma), and DDM (D310S, Calibre Scientific). Samples were divided for use in total intracellular protein and phosphorylation quantification. Total intracellular protein quantification was performed using the Micro BCA Protein Assay Kit (23235, Thermo) according to manufacturer instructions. Sample protein concentrations were interpolated from a standard curve using GraphPad Prism (v. 10.6.0). Comparison of multiple groups was performed by using one-way analysis of variance (ANOVA) with Dunnett correction. Phosphorylation quantification was performed via Western Blot by detecting phosphorylated serine and threonine residues with tyrosine, tryptophan, or phenylalanine at the -1 position or phenylalanine at the +1 position. Samples were resolved by SDS-PAGE, with an equal proportion of cell lysate from each sample loaded in each lane. Phosphorylation levels were analyzed by immunoblotting using a 1:1000 dilution of Phospho-(Ser/Thr) Phe Antibody (9631, Cell Signaling Technology) for primary staining and a 1:10000 dilution of IRDye 680RD Goat Anti-Rabbit IgG (926-68071, LICORbio). The blotted membrane was imaged using a LICOR Odyssey imaging system. Signal intensity was quantified using ImageJ 1.54 and plotted in GraphPad Prism (v. 10.6.0). Comparison of multiple groups was performed by using one-way analysis of variance (ANOVA) with Dunnett correction.

**Cell fixation for Raman spectroscopy**

Upon confluency, media was aspirated and the cells were washed with 5 mL of PBS. After removal of media, 3 mL of 0.25% trypsin-EDTA was added to the culture flask and incubated at 37°C at 5% CO2 for 5-7 min. After incubation, RPMI + 10% FBS was added to quench the trypsin reaction, and the cell suspension was collected into a 15 mL centrifuge tube. The tube was spun for 5 minutes at 400 g, and the supernatant was aspirated. Next, 1 mL of 4% paraformaldehyde was added, and the solution was mixed gently and incubated at room temperature for 10 minutes. After incubation, the tube was centrifuged for 3 minutes at 400 g at 4oC before supernatant removal. The cell pellet was resuspended with an additional 1 mL PBS and centrifuged for 3 minutes at 400 g. The supernatant was aspirated, 50 uL of PBS was added, and the solution was gently mixed. Based on cell concentration, 1-5 uL of cell solution was drop casted on a gold-coated slide (~195 nm Au, 5 nm Ti, 1 mm glass) and air dried for 5-10 minutes prior to Raman acquisition. For live cell preparation, cells were removed prior to paraformaldehyde and suspended in 50 uL of PBS.

**Targeted inhibitors for Raman spectroscopy**

Powders of high purity (>99.9%) bemcentinib, cabozantinib, and dabrafenib were each thoroughly mixed with 3 uL of PBS with a pipette. 2 uL of mixture was dropcast on a gold-coated slide and air dried for 5-10 minutes prior to Raman acquisition.

**Raman data acquisition**

Raman spectra on the dried samples were acquired using a confocal Raman spectrometer (Horiba LabRAM, Longjumeau, France) at 532 nm excitation with a 100x 0.6 NA objective at five second integration times. Using a 600 gr/mm grating, the spectral acquisition range acquired was 500-2000 $cm^{-1}$. All spectra, unless otherwise stated, were acquired with 100% power output of ~5.4 mW with a nominal spot size of ~2.1 µm. Spectra for murine sample B16-F10 and human samples SK-MEL-30 (control) and PAT-73 (bemcentinib) were collected at both 100% and 50% power using an integrated ND filter. Spectra for murine samples YUMM (cabozantinib, dabrafenib, nivolumab) were collected at both 25% and 50% power. Spectra were acquired using two approaches: a) via a rectangular grid mapping with 20 µm spacings between point measurements for throughput, and b) via targeted point-by-point selection per window region on visually identified cells to ensure approximate single-cell acquisition and minimize cell measurement duplicates. While the confocal configuration provided subcellular lateral resolution, finite axial depth of field combined with occasional 2-3 cell layering meant some grid-mapped spectra contained contributions from multiple cells. Point-by-point measurements on isolated cells ensured single-cell data. Spectra for targeted inhibitors were acquired with 25% power for dabrafenib and bemcentinib and at 100% power for cabozantinib. In total, 8,095 cell spectra were utilized for murine TIME cell differentiation, 3,594 for macrophage polarization, 9,937 for murine drug response, 33,921 for human drug response, and 23,994 for melanoma resistance determination. For biochemical validation, 2,188 cell spectra were utilized.

**Raman data acquisition for live vs fixed comparison**

Raman spectra on fixed cells and live cells at 532 nm excitation followed the preparation and measurement protocols described prior. Live cells suspended in PBS medium were placed in a liquid well constructed of a gold-coated glass slide base layer with hole-punched, double sided adhesive tape serving as the side walls. Vacuum grease was utilized on the edge of the double sided tape to prevent sample leakage, and the well was covered with a borosilicate glass cover slip to prevent sample evaporation. 4 µL of cell solution was placed in the liquid well. Spectra at 785 nm excitation was acquired using a confocal Raman spectrometer (Horiba XploRA+, Longjumeau, France) using a 50x/0.65 NA objective (Olympus LCPLAN 50X IR) with a 0.5 correction collar. Using a 1200 gr/mm grating, the spectral acquisition range acquired was 500-2000 $cm^{-1}$. All spectra were acquired with 100% power output of 26.34 mW with a nominal spot size of ~1.5 μm. Spectra were acquired at 20 second integration times. The 50x objective was additionally used for sequential time series acquisitions to minimize background contribution from the cover slip.

**Data preprocessing**

Spectra were first filtered by an intensity count threshold to remove detector saturated measurements. Unprocessed Raman spectra were then despiked using a modified Whitaker-Hayes method to remove cosmic rays and spiked pixel regions on the CCD[1]. Upon despiking, spectra were denoised via wavelet thresholding. The python package skimage was used to perform our smoothing, and BayesShrink was chosen as our wavelet coefficient threshold selection method[2]. Following our spectral smoothing, a baseline correction was applied using a combination of polynomial fitting (4th order) and adaptive iteratively reweighted Penalized Least Squares (airPLS)[3]. The correction method was implemented using the python packages PeakUtils and BaselineRemoval[4,5]. Each baseline-subtracted spectrum was internally normalized by replacing the intensity of each wavenumber with the standard score, calculated from the mean and standard deviation of the intensities in the spectrum, to enable cross-spectrum comparisons. The biological fingerprint region of 600-1800 $cm^{-1}$ of the processed spectra was isolated for analysis. Off-cell measurements from grid mapping, stage translation drifts, and background media and poor-quality spectra characterized by high background to cell signal intensities were filtered utilizing a combination of one-class support vector machines and major biological bands z-scores for all datasets. The set of one-class support vector machines (SVMs, sklearn)[6] was trained using spectra from relevant backgrounds (gold slide, phosphate-buffered saline, supernatant, polystyrene).

### Supplementary Methods: Data Analysis

#### Spectral clustering

Preprocessed data were subjected to linear dimensionality reduction by PCA (sklearn) to reduce the feature size to a set of principal components explaining ≥95% of the dataset variance. The transformed dataset aided in visualization and in certain cases, improved classification runtimes. Manifold learning (UMAP, umap and Scanpy)[7,8] was applied to the PCA-reduced dataset to visualize the samples in a two-dimensional embedding, and hyperparameters random_seed=42, n_neighbors=25, min_dist=0.5, spread=1, n_epoch=500 were used across all datasets, with adjustments of the following: min_dist=0.3, spread=2 for murine cell differentiation described in Figure 2b; n_neighbors=30 for mice drug response, min_dist=0.3 for human drug response, and min_dist=0.3, spread=3 for patient data. Effects of hyperparameters on UMAP embedding are highlighted in Supplementary Figure 17. Leiden clustering[9] (Scanpy) with resolution 0.3 (human response) and 0.5 (patient dataset) was used to examine spectral differences among clusters. Heatmap (differential Raman) analysis was performed by first shifting our dataset by a constant c, where c is the value needed to raise our minimum value to nonzero. After, the median value of the wavenumber features were determined from our control group. Median was selected over arithmetic mean for improved robustness against dissimilar group distributions[10]. In the case of the drug screening assay, the untreated cells served as control. Finally the fold change was determined for each sample through the log2 of the ratio between sample features and the control group. Log2 was chosen such that a two fold increase (doubling of quantity) resulted in a log2 fold change of +1 whereas a two fold decrease (halving of quantity) resulted in a log2 fold change of -1. Wavenumber selections for the differential intensity heatmaps were either determined from the top wavenumbers affecting classification accuracy by perturbation calculations (see feature importance section below) or by the combination of most prominent peak selection in the average spectra (SciPy)[11] and highest variant wavenumbers using Wilcoxon rank-sum (Scanpy). Cell spectra were subsampled for heatmap visualization.

#### Spectral barcodes

Barcodes using the top 50 variant wavenumbers were generated from averaging the log2 fold change of 500 randomly selected spectra from each sample. Cell spectra with predicted probabilities of >0.8 and <0.2 from each of the drug determination classifiers were selected for high-likelihood resistant and sensitive barcodes, respectively. Barcodes were color bound symmetrically with a ±0.5 for bemcentinib and cabozantinib, ±0.75 for nivolumab, and ±1 for dabrafenib and nivolumab + relatlimab to maximize treatment comparability and reduce individual patient outliers and label imbalance associated with our limited patient dataset.

#### Leiden cluster cell response labeling

For each Leiden cluster, the number of cells labeled resistant and sensitive was determined by examining the clinical and cell viability data. Cluster population proportions were then calculated relative to the total number of cell spectra assigned to the cluster, where the sum of resistant + sensitive + unlabeled cell spectra proportions = 1.

**Feature importance**

Raman wavenumbers importance were determined by analyzing the classification accuracy dropoff from spectra masking across all wavenumbers[12]. First, a 10-split stratified shuffle cross-validation (sklearn) was utilized with an 80:20 training:test split. Utilizing the random forest model discussed prior, we then determine the general classification performance as each wavenumber feature is perturbed with a Voigt profile of alpha=5, gamma=2 of the pre-transformed data for each test split. The average classification accuracy and standard deviation was then calculated for each feature. The Raman bands with the biggest effect on classification were selected by a descending sort of the largest peaks (SciPy) associated with 1-classification accuracy.

**Construction and implementation of resistance determination model**

A random forest multiclass classifier (sklearn) was constructed for each drug of interest using hyperparameters max_depth = 20, max_features = "sqrt", n_estimators = 275. Hyperparameters were selected by grid searching on the nivolumab training dataset and utilized for remaining determination models. Additionally, class_weight="balanced" was used to account for imbalanced training datasets. Commercial human cell lines and patient samples without additional treatments were used for our training and test datasets. We augmented the inhibitor training set with commercial cell line spectra. Each training patient sample was required to have 500+ spectra. Labels 0 (sensitive) and 1 (resistant) were annotated based on clinical observations and in vitro cell viability assessments. Cell viability cutoffs were set at 70% (targeted inhibitors) and 60% (immunotherapy) with edge cases (±2%) annotated sensitive. In summation, the first passages of patient samples PAT-52, PAT-54, PAT-63, PAT-73, PAT-93 and cell lines A375, SK-MEL-24, and SK-MEL-30 were utilized in our training set. Class probabilities from 0 to 1 as defined by class probabilities in our RF model were determined for each spectra, where values closer to 0 and 1 corresponded with more sensitive and more resistant, respectively. A binary label of resistant or sensitive could then be assigned based on a desired determination cutoff threshold for each drug protocol of interest. In our standard model, we chose 0.5 as our determination cutoff for the aggregated patient response. Labeling assignments for each treatment model were also calculated for determination cutoff ranges from 0 to 1 at 0.1 intervals to select optimized cutoffs. For each patient, 25,000 mini patients were generated by grouping 100 cell spectra randomly selected using uniform weighting from the patient superset. Drug resistance probability was predicted for each mini patient spectra using our pre-trained RFs, and the quantitative response of a mini patient was calculated by the mean of the spectra likelihoods. The patient response determination was then assessed by examining the median, 5th, and 95th percentile.

**scRNA-seq analysis**

Data were aligned with cellranger 9.0.0 to reference genome GRCh38-2024-A[13]. Data were imported into python as anndata objects for further analysis using the Scanpy[8,14]. Preprocessing was performed on each sample individually. Cells with fewer than 100 unique genes or more than 10% mitochondrial counts were removed. Doublets were removed using the DoubletDetection package with Louvain clustering, 20 iterations, 1e-16 p-value threshold, and 0.5 voter threshold[15]. Integration was performed with scVI using the top 2000 highly variable genes[16–18]. The model was trained for 100 epochs. Leiden clustering was performed at resolution 0.9. Differential expression analysis was performed using the scVI differential expression module, and cells were manually annotated as tumor cells and fibroblast-like cells using canonical marker genes. One cluster that had a mixed gene expression profile was marked as unknown. The scANVI framework was then used for further integration using the scVI annotations as initial labels[19]. Leiden clustering was performed at resolution 1.0. Differential expression analysis was performed using the scVI differential expression module, and cells were manually annotated as tumor cells and fibroblast-like cells using canonical marker genes.

**Statistical Analysis on Raman spectroscopy**

Statistical analysis was performed at individual wavenumbers of interest, selected based on prior literature and preliminary spectroscopic evaluation. For each center wavenumber, intensities were integrated over ± 5 $cm^{-1}$ window to account for spectral variability. Two complementary statistical approaches were employed to account for the hierarchical structure of the data (cells nested within biological replicates) and to provide robust assessment of treatment effects. Single-cell measurements (μ=243 cells, n=3 biological replicates per treatment) were analyzed using 1) one-way analysis of variance (ANOVA) with Tukey's HSD (scipy, statsmodels)[20] on biological replicate-averaged integrated intensities, and 2) linear mixed-effect models (LMMs) on single-cell data with biological replicate as random effect. LMMs were fitted using the Powell optimization method to ensure convergence. LMM p-values were adjusted for multiple testing using the Benjamini-Hochberg false discovery rate (FDR) procedure across all tests ($\alpha = 0.05$). Random effects variance was minimal, indicating high between-replicate consistency. Both approaches are reported for transparency; statistical significance defined as $p<0.05$.

**Statistical analysis on intracellular lipid, protein, and phosphorylation quantification**

Lipid: Comparison of multiple groups was performed by using one-way analysis of variance (ANOVA) with Dunnett correction. Protein: Sample protein concentrations were interpolated from a standard curve using GraphPad Prism (v. 10.6.0). Comparison of multiple groups was performed by using one-way ANOVA with Dunnett correction. Phosphorylation: Signal intensity was quantified using ImageJ 1.54 and plotted in GraphPad Prism (v. 10.6.0). Comparison of multiple groups was performed by using one-way analysis of variance (ANOVA) with Dunnett correction.

### Supplementary Figures

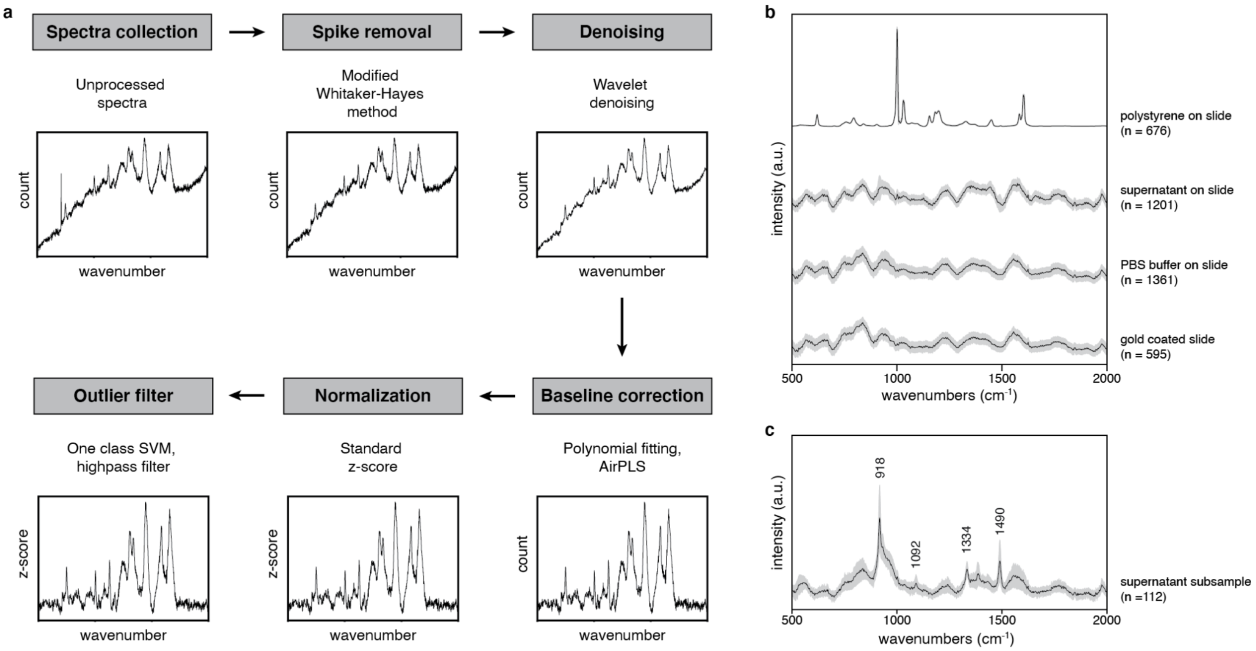


**Supplementary Figure 1. Workflow of Raman spectra preprocessing.** A) Representative workflow of the preprocessing steps taken for each spectrum. Preprocessing was essential to normalize and remove artifacts for cell spectra comparisons. An example spectrum of RAW264.7 is shown, highlighting the post-processed spectra after each step. B) Averaged spectra with ±1 standard deviation of post-processed background and outlier spectra acquired for our automated data filter step. The data filter step utilizes a combination of single class support vector machines (SVMs) and high-pass z-score values, and occurs after the preprocessing workflow highlighted in (A). C) A subset of spectra in the supernatant spectral set in (B) where major peak assignments are shared with polymers such as polyoxymethylene copolymer (POM-C) found in literature.

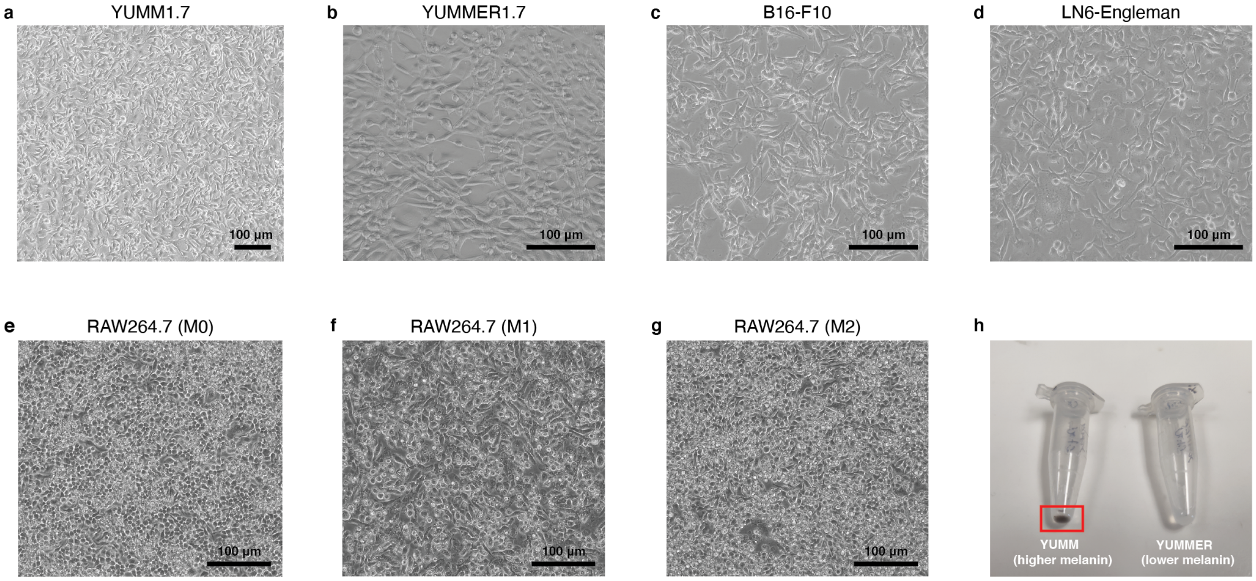


**Supplementary Figure 2. Bright field images of mouse cells.** A-G) Brightfield images of live murine cell lines. Images highlight morphological differences associated across cell types in cancer and immune cells and functional phenotypes in polarization states. Scale bar, 100 µm. H) Image of YUMM1.7 and YUMMER1.7 cell lines highlighting contrast in melanin pigmentation (higher melanin in the left tube highlighted with a red box), which was captured in our Raman analysis.

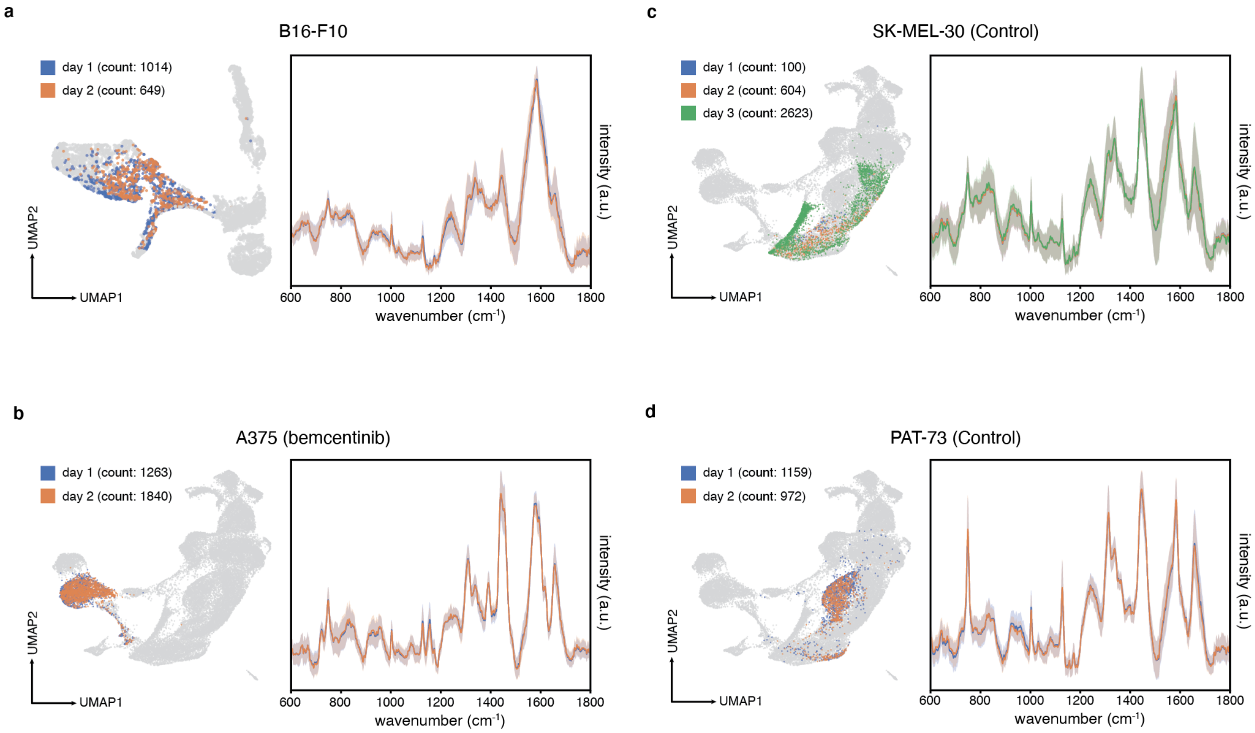


**Supplementary Figure 3. Day to day variations in Raman spectra acquisition.** Select mouse cell lines and human patient-drug treatment samples were measured across multiple days to examine spectra variability relating to sample preservation, environmental fluctuations, and instrumentation inconsistencies for (a) B16-F10, (b) A375-bemcentinib, (c) SK-MEL-30-control, and (d) PAT-73-control. Findings show consistent overlays across our embedded space (left) for a given sample-treatment protocol, and average spectra well within the ±1 standard deviation (right).

a

b

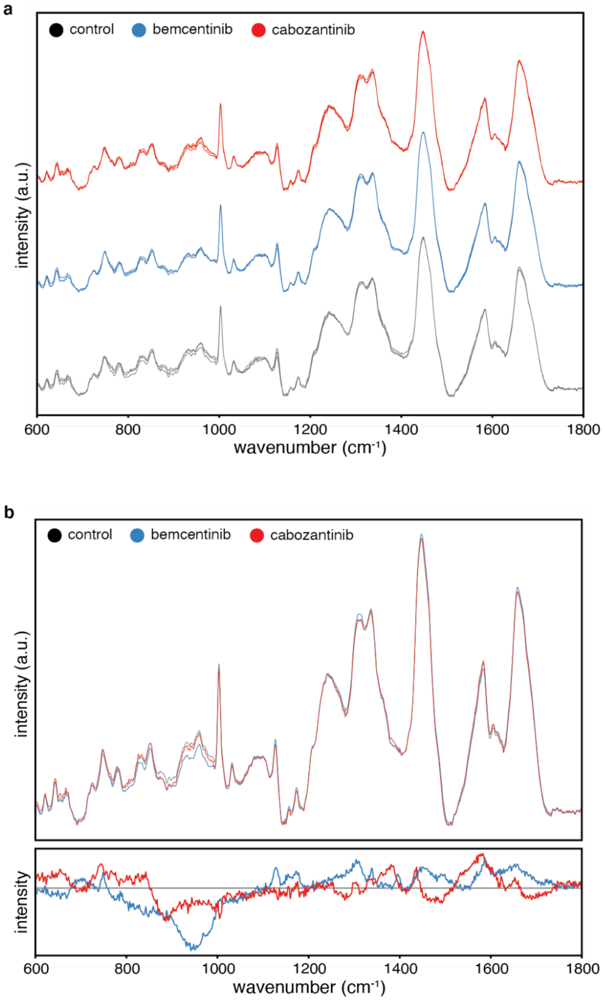


c

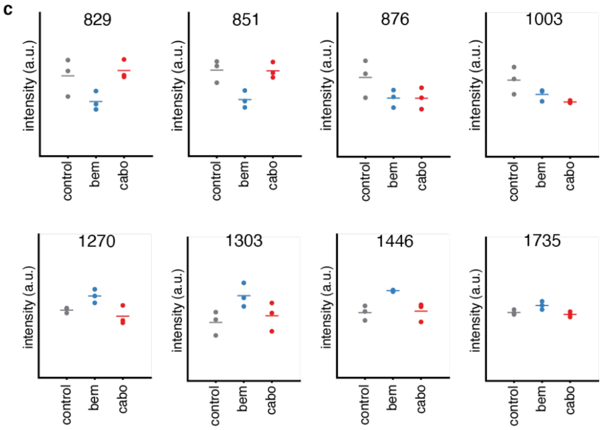


d

| wavenumber (cm-1) | 1-way ANOVA | Post-hoc | | Linear Mixed-Effects (FDR-adj) | |
|---|---|---|---|---|---|
| | | Tukey (bem) | Tukey (cabo) | LMM-FDR (bem) | LMM-FDR (cabo) |
| 829 | 0.060 | 0.122 | 0.878 | 0.055 | 0.673 |
| 851 | **0.013** | **0.019** | 0.996 | **0.004** | 0.993 |
| 876 | 0.173 | 0.222 | 0.220 | 0.090 | 0.090 |
| 1003 | **0.053** | 0.185 | **0.048** | 0.071 | **0.010** |
| 1270 | **0.029** | 0.103 | 0.550 | **0.029** | 0.359 |
| 1303 | 0.096 | 0.095 | 0.808 | **0.029** | 0.671 |
| 1446 | **0.014** | **0.019** | 0.966 | **0.005** | 0.869 |
| 1735 | **0.031** | 0.078 | 0.762 | **0.029** | 0.600 |

e

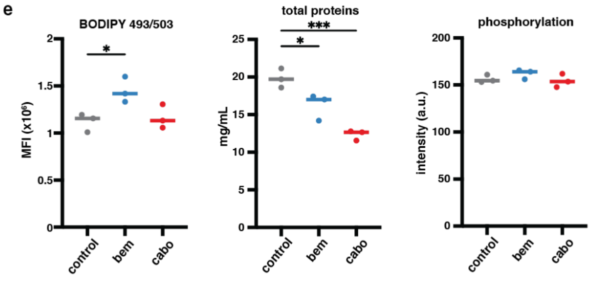


**Supplementary Figure 4. Lipid and protein quantification of post-treatment YUMMER1.7 cells.** A) Mean Raman spectra (μ=243 cells) of control (black), bemcentinib-treated (blue), and cabozantinib-treated (red) YUMMER1.7 cells (n=3 biological replicates) in close agreement within treatment groups. B) Top: Mean Raman spectra of cell response to treatment groups (μ=729 cells). Bottom: Spectral differences of post-treated samples relative to untreated (control). C) Intensity distributions at representative wavenumbers showing post-treatment differences. Lipid-only (1735 $cm^{-1}$) and lipid-dominant (1270, 1303, 1446 $cm^{-1}$) peaks commonly used in composition analysis showing post-bemcentinib spectra with increased intensities and post-cabozantinib spectra with relatively same intensity to control[21–26]. Protein only peaks (1003, 876 $cm^{-1}$) showing intensity decreases across post-bemcentinib and post-cabozantinib spectra, with a larger decrease on average for post-cabozantinib[22,24,27,28]. Tyrosine-related peaks known as fermi resonance bands (829, 851 $cm^{-1}$) showing an intensity decrease in post-bemcentinib, with no doublet collapse or change in tyrosine-associated 1205 $cm^{-1}$ intensities likely indicating reduced phosphorylation-related changes[29–31]. D) Statistical analysis of select Raman spectral peaks. One-way ANOVA tested overall differences among protocols. Post-hoc pairwise comparisons performed using Tukey's HSD (correcting for three comparisons) and linear mixed-effect models (LMM) with false discovery rate (FDR) correction across all tests (16 tests, Benjamini-Hochberg procedure). Bold values indicate $p<0.05$. See Supplementary Table 4 for Raman peak assignments. E) Cell lipid and protein quantification. Left: Intracellular neutral lipid content by BODIPY 493/503 flow cytometry showing post-bemcentinib cells having elevated lipid content. Middle: Total protein quantification by BCA assay reveals decreased protein in both post-bemcentinib and post-cabozantinb persistent cells. Right: Western blot analysis using a phospho-motif antibody showing no significant differences among treatment groups.

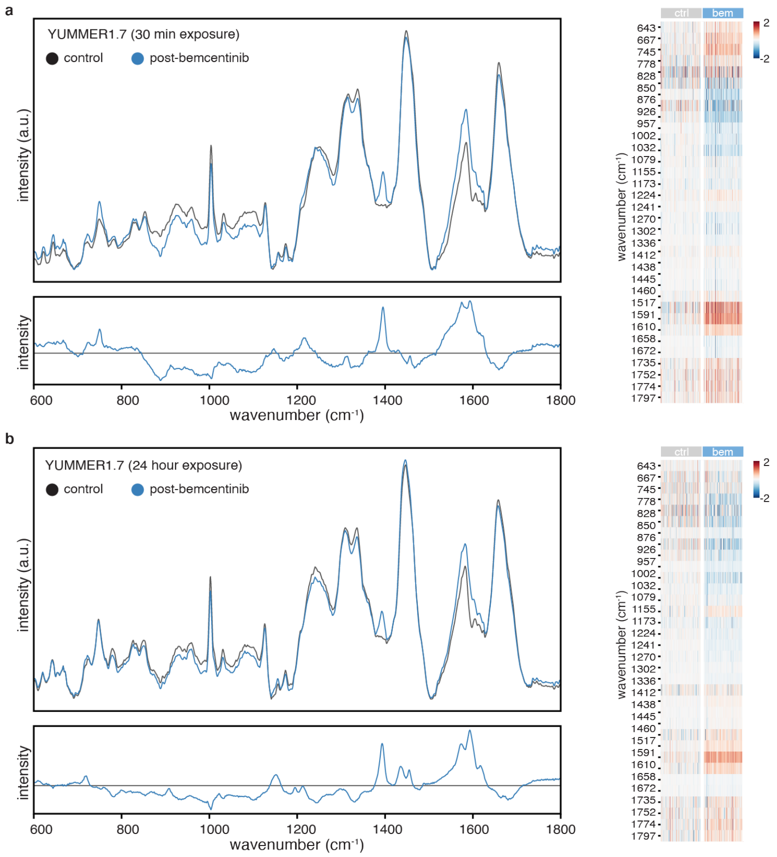


**Supplementary Figure 5. Temporal dynamics of Raman spectral changes in bemcentinib-treated YUMMER1.7 cells.** YUMMER1.7 cells were exposed to bemcentinib for 30 minutes (a) or 24 hours (b). Left panels: Mean normalized Raman spectra of control (black) and post-bemcentinib (blue) cells (top) with corresponding difference spectra (bottom, post-treatment minus control). Right panels: Differential Raman intensities across select wavenumbers, with cells grouped by treatment conditions. Color scale represents fold change from -2 to +2. Pronounced spectral changes were observed at 30 minutes followed by partial recovery towards baseline at 24 hours across lipid (958, 1447, 1658 $cm^{-1}$), protein (750, 930, 958, 1003, 1030, 1241, 1447, 1658 $cm^{-1}$), and nucleic acid (750, 1241, 1337 $cm^{-1}$) peaks. This temporal trajectory is consistent with initial acute cellular stress response followed by adaptive remodeling.[32–36] Temporal trajectories at nucleic acid-associated bands (780-800 $cm^{-1}$) differed from the intensity changes characteristic of apoptotic cells, supporting an adaptive remodeling interpretation.[37–40] Spectral peaks reflective of remnant bemcentinib which was seen in prior analysis (Supplementary Figure 6) demonstrate Raman's high sensitivity.

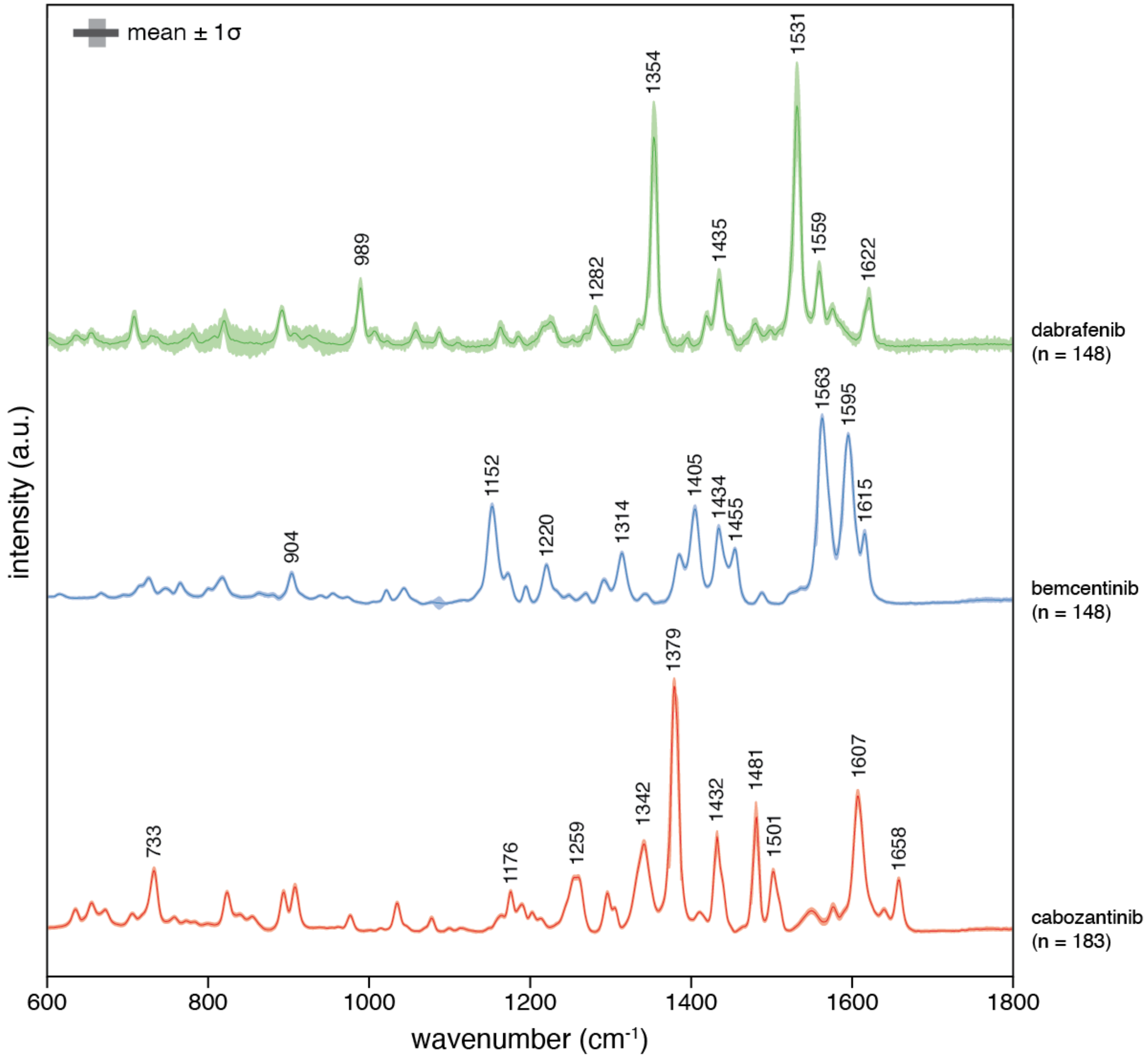


**Supplementary Figure 6. Raman spectra of molecular inhibitors.** Mean normalized spectra and ±1 standard deviation for targeted inhibitors cabozantinib, bemcentinib, and dabrafenib. Wavenumber assignments are annotated for each averaged spectra.

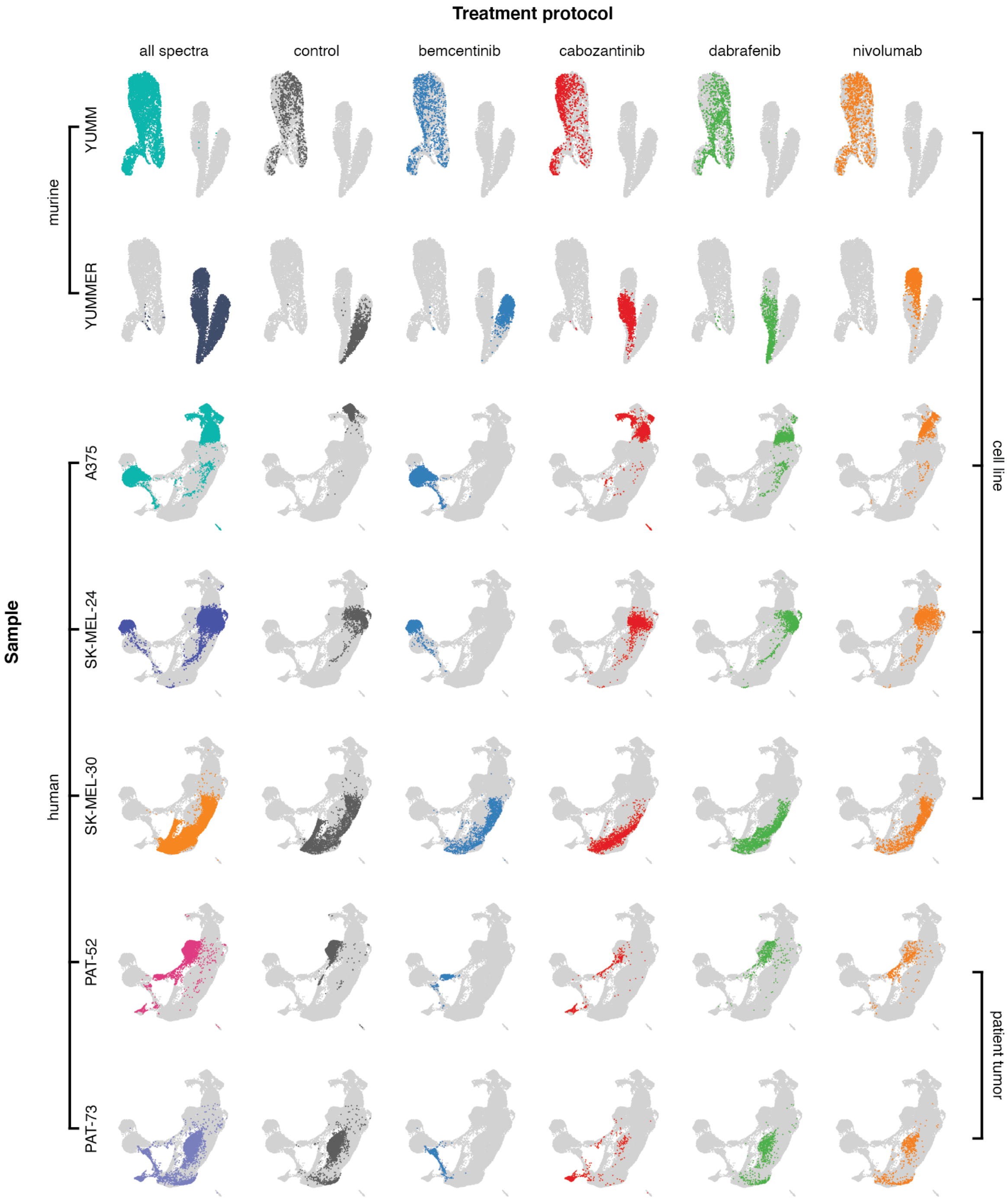


**Supplementary Figure 7. One-vs-all cell spectra for mouse and human samples.** Latent space localizations of mouse and human line samples discussed in Figure 3 and 4. For each sample, spectra are highlighted with respect to cell procurement source (row) and treatment protocol (column). The first column encompasses all affiliated cell spectra acquired for a given sample, and each successive column represents either an untreated cell set (control) or persistent cells measured following a treatment protocol.

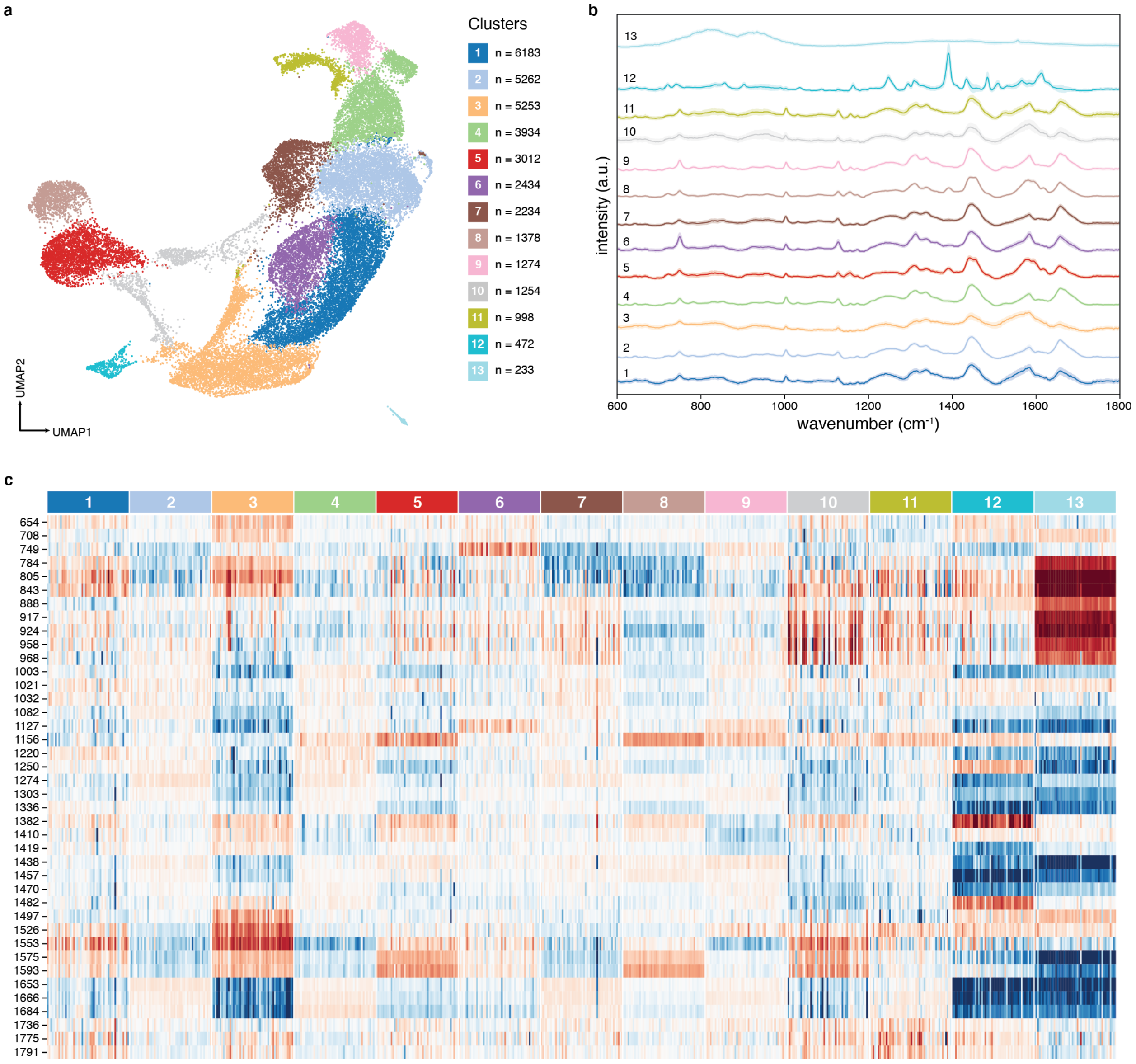


**Supplementary Figure 8. Leiden clustering of human samples.** A) UMAP embedding of Raman profiles from human melanoma response dataset grouped by Leiden clustering. Hyperparameters selected were nearest neighbors = 25, minimum distance = 0.3, number of epochs = 500, and Leiden resolution = 0.3. B) Mean normalized spectra and ±1 standard deviation across each Leiden grouping. Note, cluster 13 shows typical spectra from our substrate, while cluster 12 shows major bands found in spectra observed in cabozantinib. C) Differential wavenumber intensities of the median fold change across Leiden clusters showing distinct Raman "expression" for each cluster.

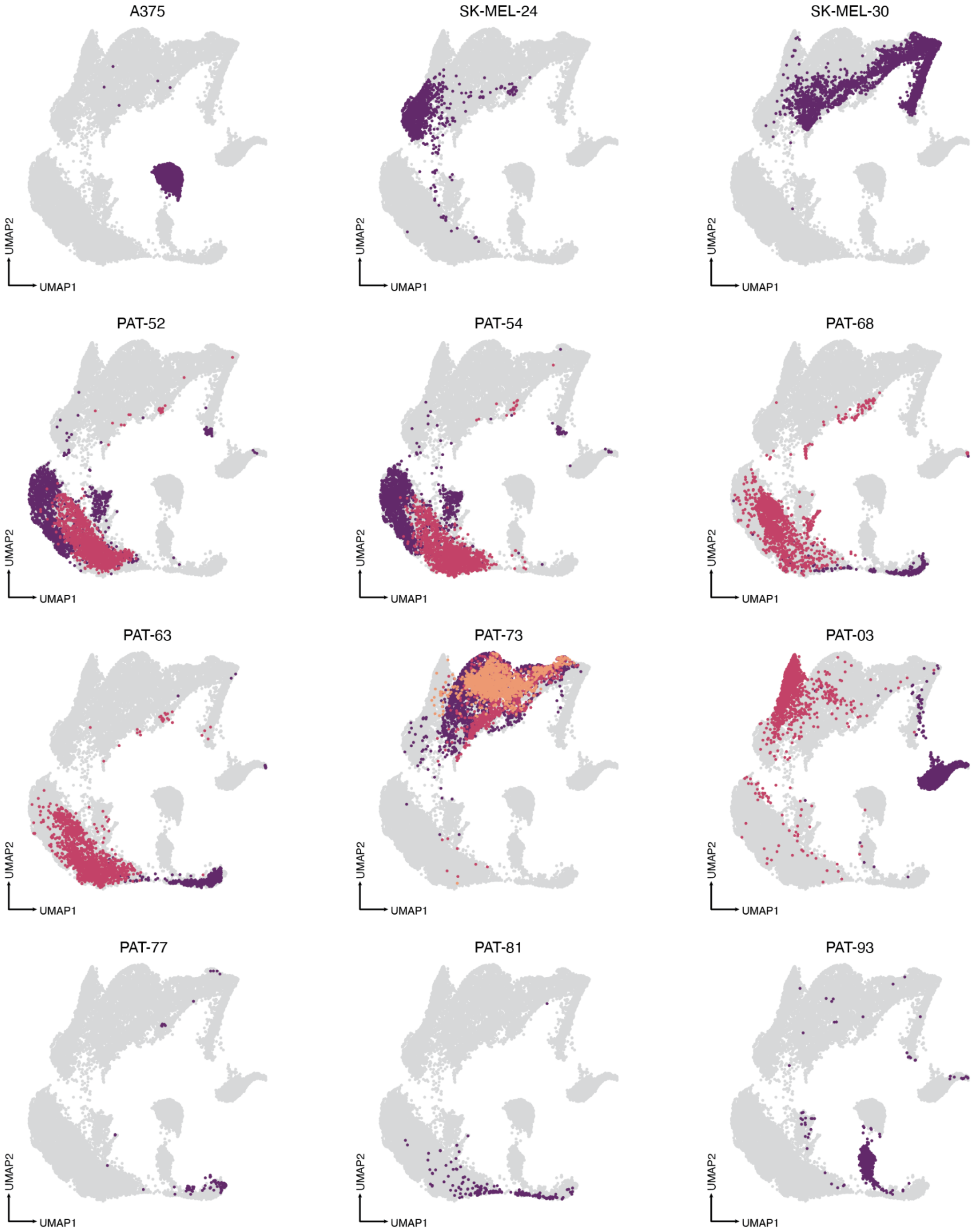


**Supplementary Figure 9. Passage to passage variations for select patient samples.** UMAP embedding of Raman profiles from patient melanoma and commercial cell lines. First passage is colored purple, second passage colored red, and third passage, if applicable, is colored orange. Two passages were analyzed for all patient samples above, except for PAT-73 in which three passages were analyzed. Commercial cell lines are represented in the first row, primary tumors in the second, and immunotherapy-refractive patients in the remaining rows.

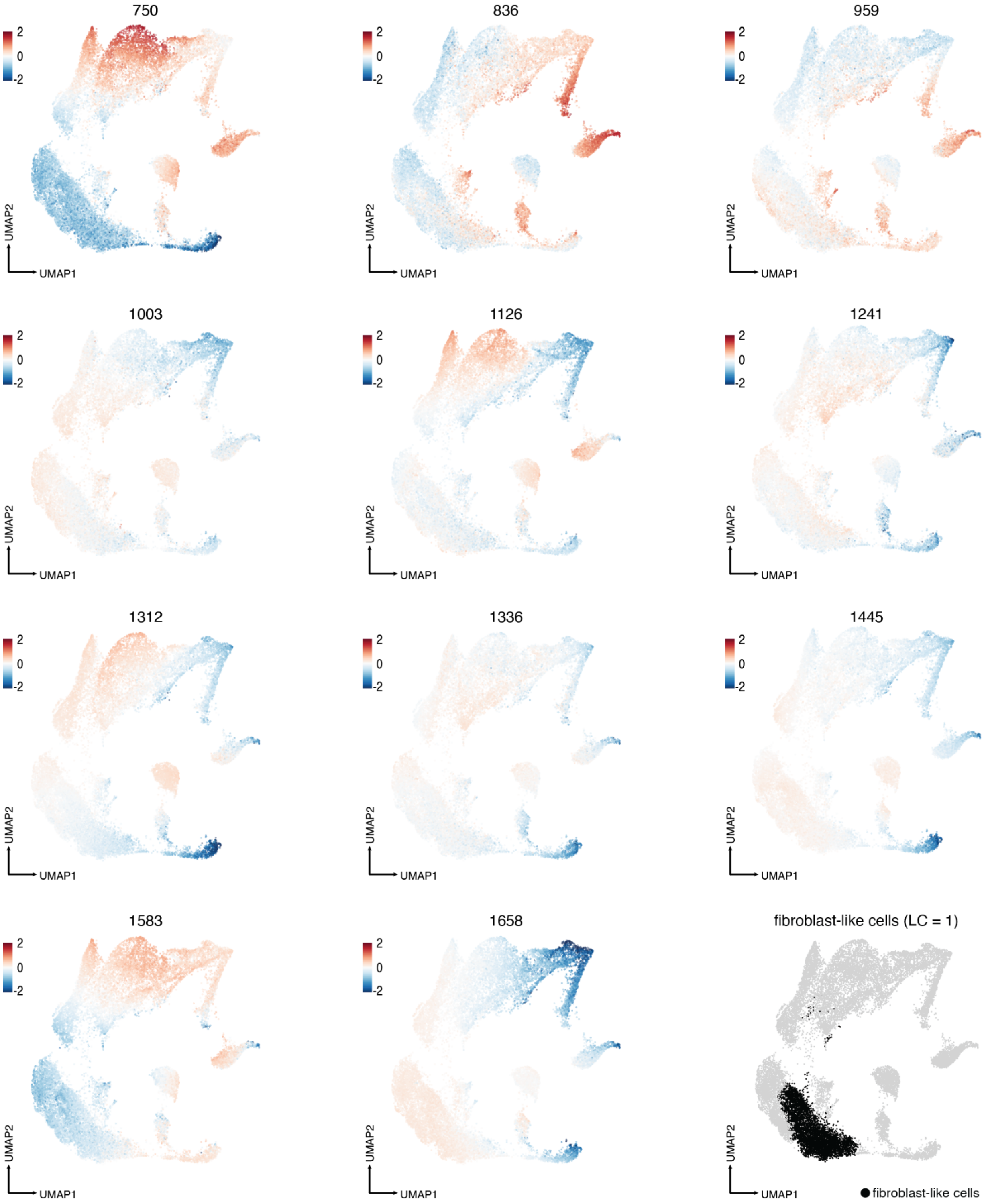


**Supplementary Figure 10. Raman intensity fold change levels for patient samples by UMAP.** Median fold change across select highly variant wavenumbers for our patient dataset, showing elevated or reduced "expression" across localized regions in latent space. Fibroblast-like cell profiles (cell spectra highlighted in black in the last subplot and corresponded to Leiden grouping 1) had elevated "expressions" across 1658 and reduced "expressions" across 750, 1312, and 1583 $cm^{-1}$.

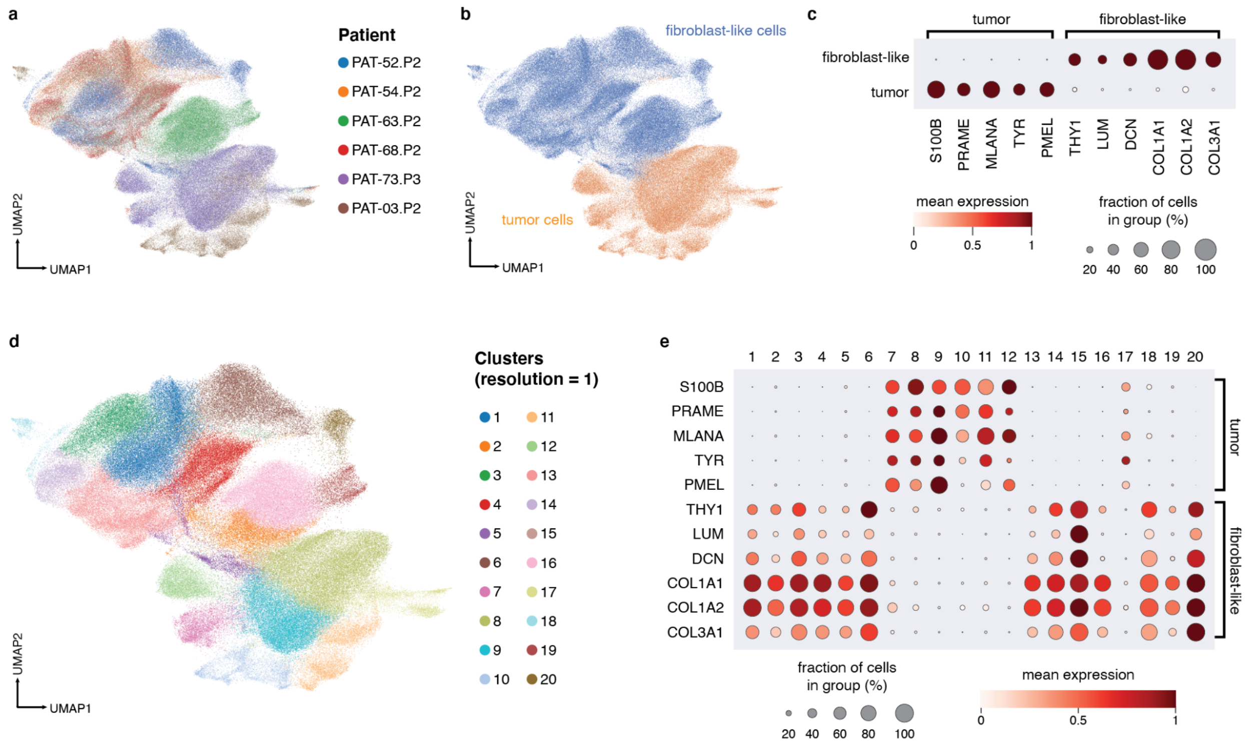


**Supplementary Figure 11. Single-cell RNA sequencing of patient melanoma samples.** Latest passages from samples PAT-52, PAT-54, PAT-63, PAT-68, PAT-73, and PAT-03 were sequenced. A) Transcriptomic UMAP colored by patient. B) UMAP colored by cell type based on canonical marker genes. C) Dotplot of marker genes associated with tumor-like and fibroblast-like features. D) Leiden clustering at resolution 1.0. E) Dotplot of marker genes grouped by Leiden cluster. We observe bimodal distribution in the expression of these markers.

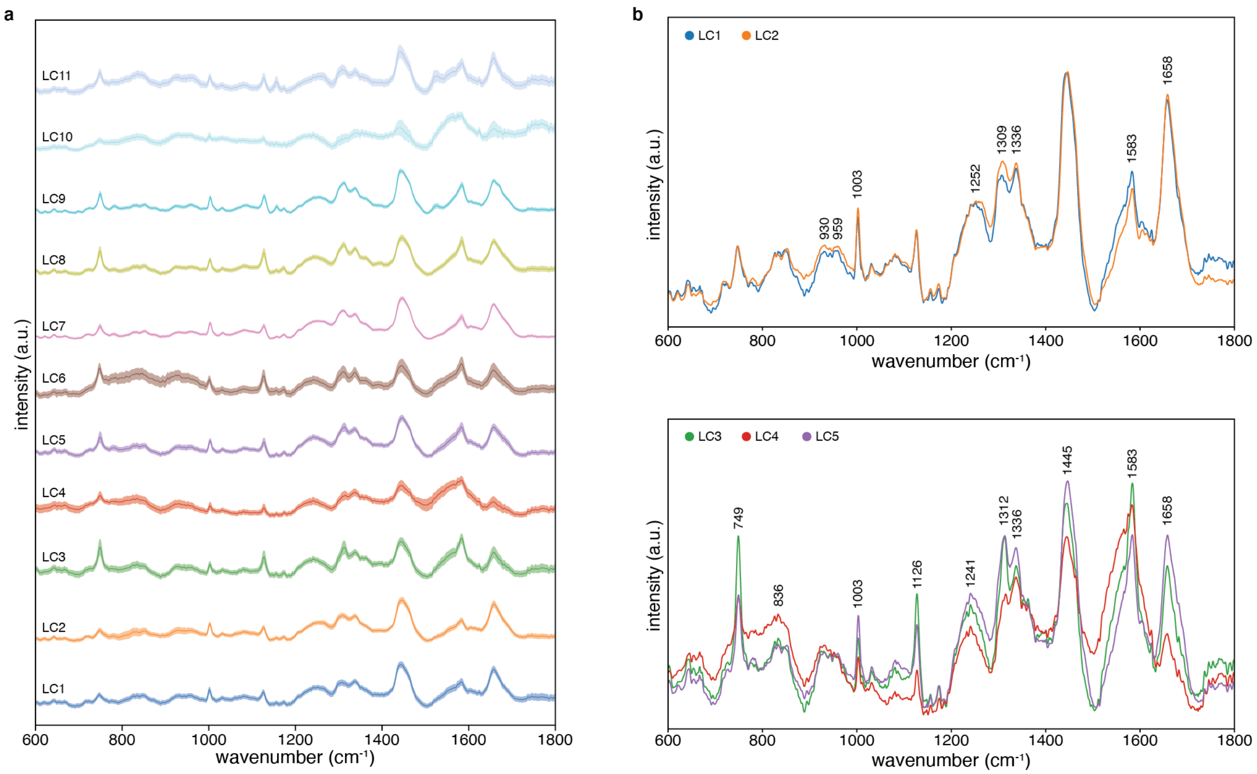


**Supplementary Figure 12. Leiden clustering for human cell lines and patient samples.** A) Mean normalized Raman spectra and corresponding ±1 standard deviation of cells residing in each Leiden cluster (resolution = 0.5) from Figure 5a. B) Spectra comparison between cluster 1 and 2, highlighting differences between fibroblast-like cells (LC1) and tumor cells (LC2). C) Spectra comparison between clusters 3-5, highlighting large phenotype heterogeneity across tumor cells.

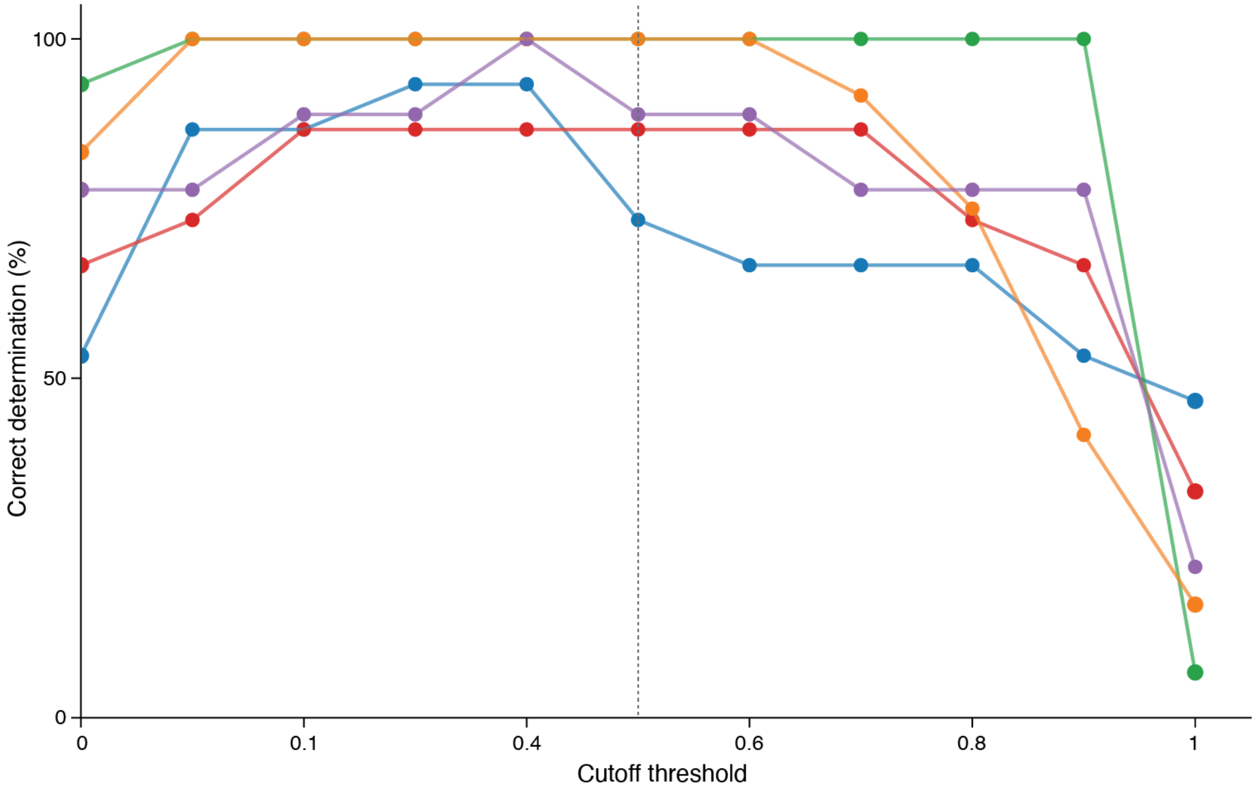


**Supplementary Figure 13. Response determination accuracy vs. threshold selection.** Examination of response determination accuracy of our prediction models across our patient samples as we vary the cutoff threshold (designating resistant and sensitive labels) of the median mini patient response. Assessment accuracy benchmarked against clinical outcomes and *in vitro* cell viability was determined for each drug classifier at 0.1 cutoff intervals. A 0.5 cutoff (dotted, grey line) was selected for all drugs of interest in our main analysis. For cabozantinib (red), dabrafenib (green), and nivolumab (orange) classifiers, the chosen cutoff threshold of 0.5 for our analysis provided the highest assessment accuracy. For bemcentinib (blue) and nivolumab + relatlimab (purple) classifiers, better assessment accuracies were found at lower cutoff values.

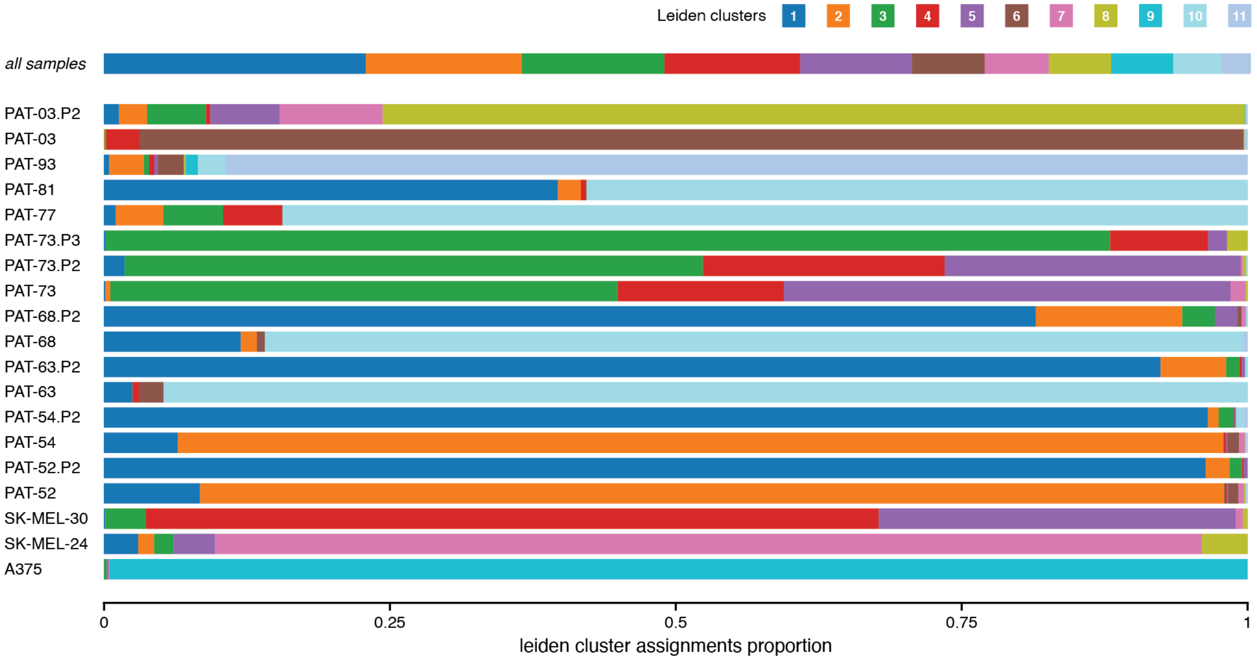


**Supplementary Figure 14. Leiden cluster designation for patient-derived samples.** Leiden cluster assignment proportions across each sample utilized in our patient prediction workflow, as well as the leiden cluster distribution encompassing our global dataset.

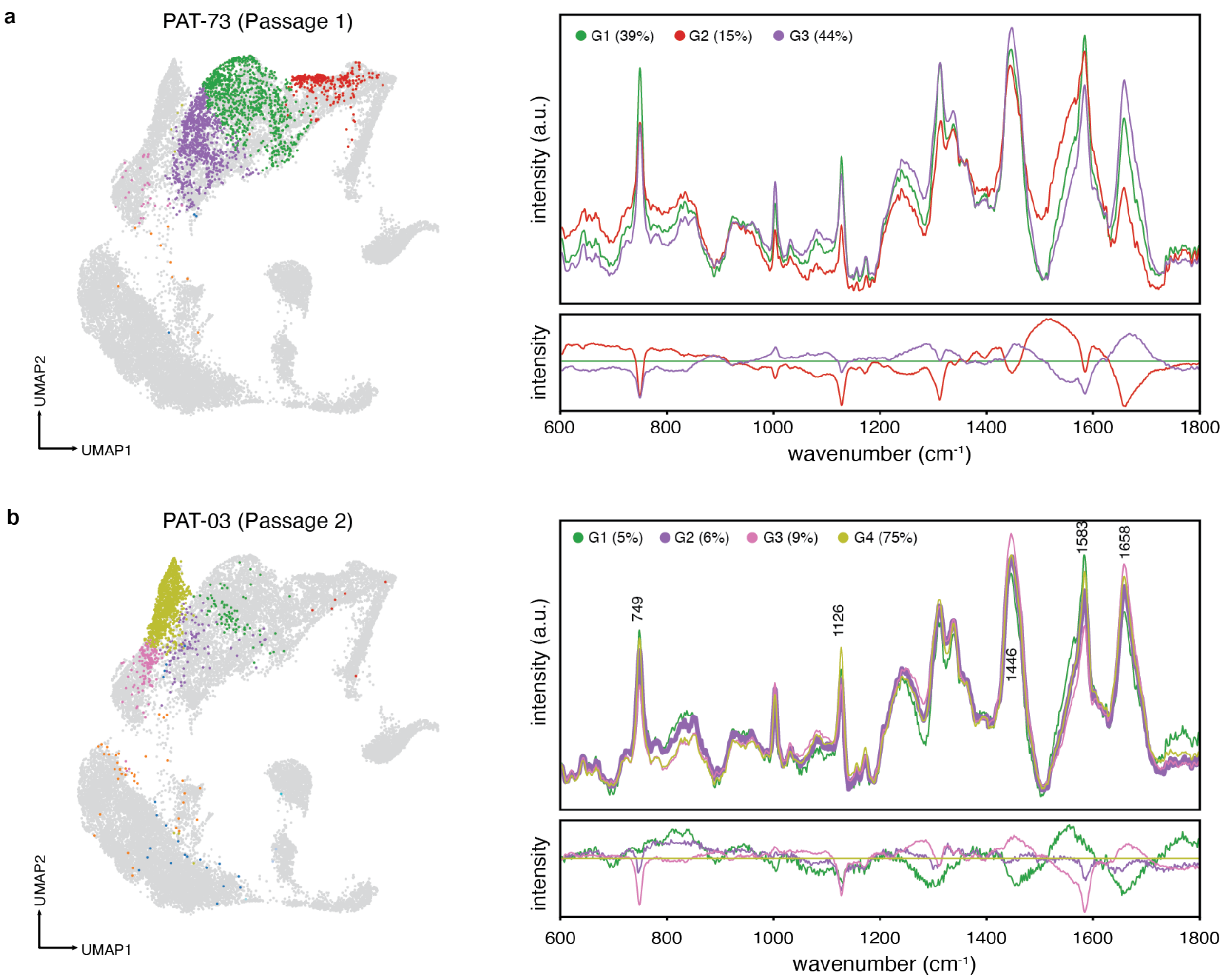


**Supplementary Figure 15. Patient samples cell subpopulations.** Select patient samples PAT-73 Passage 1 (A) and PAT-03 Passage 2 (B) showing intratumoral heterogeneity based on Leiden cluster assignments. For each section, Left: UMAP embedding of all measured cell spectra, colored by the assigned Leiden cluster found in Supplementary Figure 14. Right: Above is the mean average spectra of major groupings where a major group is defined as encompassing >5% of patient spectra, and bottom is the spectral differences among groups relative to largest grouping below. The % of patient spectra belonging to a major grouping is designated in the legend of each subfigure. Major differentiating peak assignments of (A) can be found in Supplementary Figure 12, and band assignments of 749, 1003, 1126, 1252, 1446, 1583, and 1658 $cm^{-1}$ are identified for (B). Band assignments can be found in Supplementary Table 4.

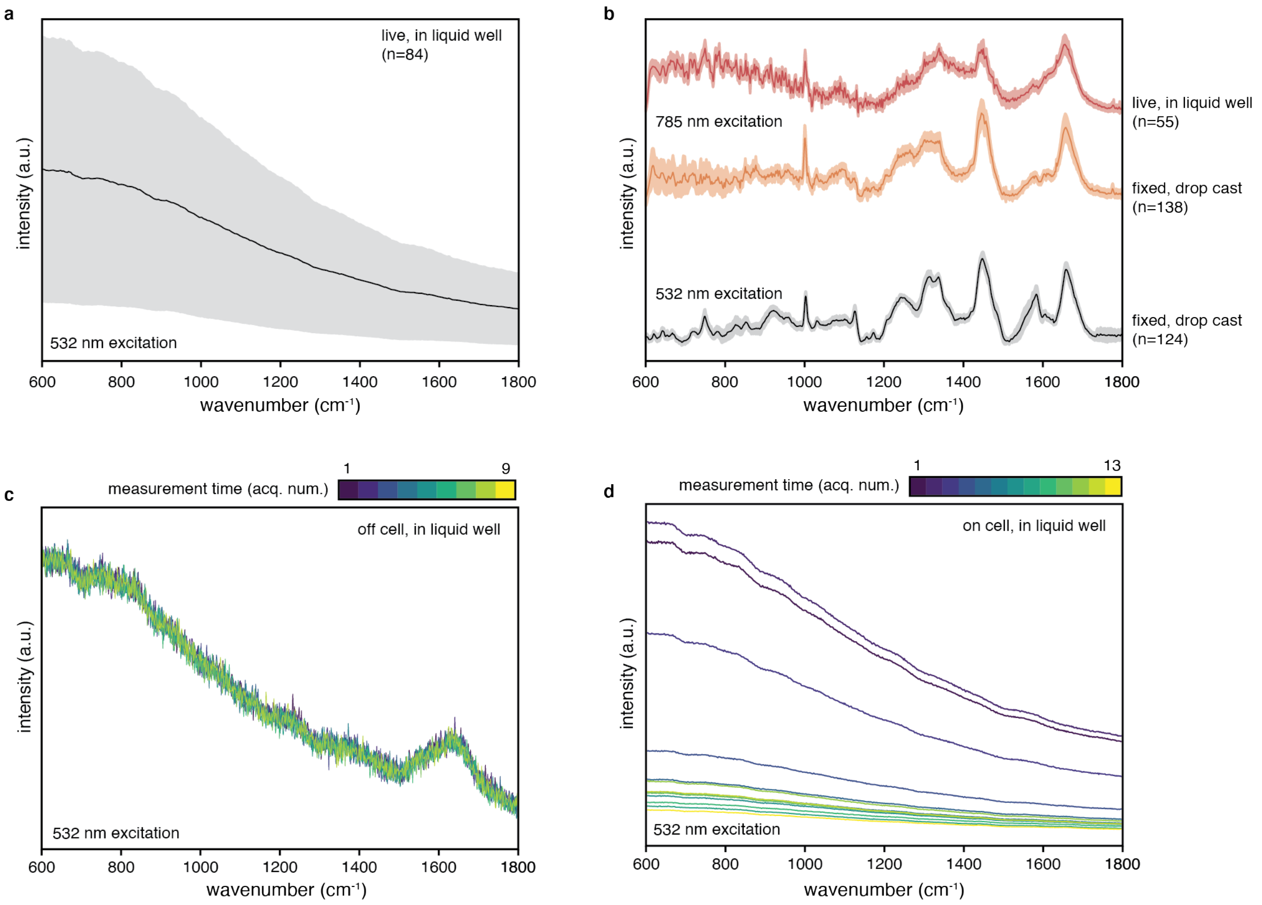


**Supplementary Figure 16. Cell spectra differences across preparation conditions.** YUMMER1.7 cells were prepared with different protocols for Raman measurements: 1) fixed and drop casted, and 2) live and in solution. A) Despiked and denoised average live cell spectra with ±1 standard deviation (n=84) under 532 nm excitation, showing high background fluorescence signal from the glass cover slip and autofluorescence of cells in PBS. Our fixed, drop-casted cell spectra exhibited weaker fluorescence signal. B) Averaged normalized spectra with ±1 standard deviation of live in suspension (n=55, red) and fixed, drop-casted (n=138, orange) cells under 785 nm excitation exhibited major spectral differences at 1000, 1250, 1312, 1341, 1447, 1580, and 1658 $cm^{-1}$, corresponding to lipid and protein bands likely arising from the presence of formaldehyde adducts, matching prior studies[41]. 785 nm excitation reduced fluorescence signal observed at shorter wavelengths. Averaged spectra with ±1 standard deviation of post-processed fixed and drop-casted cells (n=124, black) under 532 nm excitation serve as reference. Peak differences in our fixed spectra at 532 nm and 785 nm can be attributed to variations in vibration resonance enhancements that are excitation wavelength dependent. C, D) Time series (sequential) acquisitions of live in suspension samples at the same coordinate point, with each successive measurement mapped to an acquisition number. Off-cell spectra (C) showed minimal background signal, whereas on-cell spectra had high background that was reduced with successive measurements, likely from autofluorescence photobleaching[42].

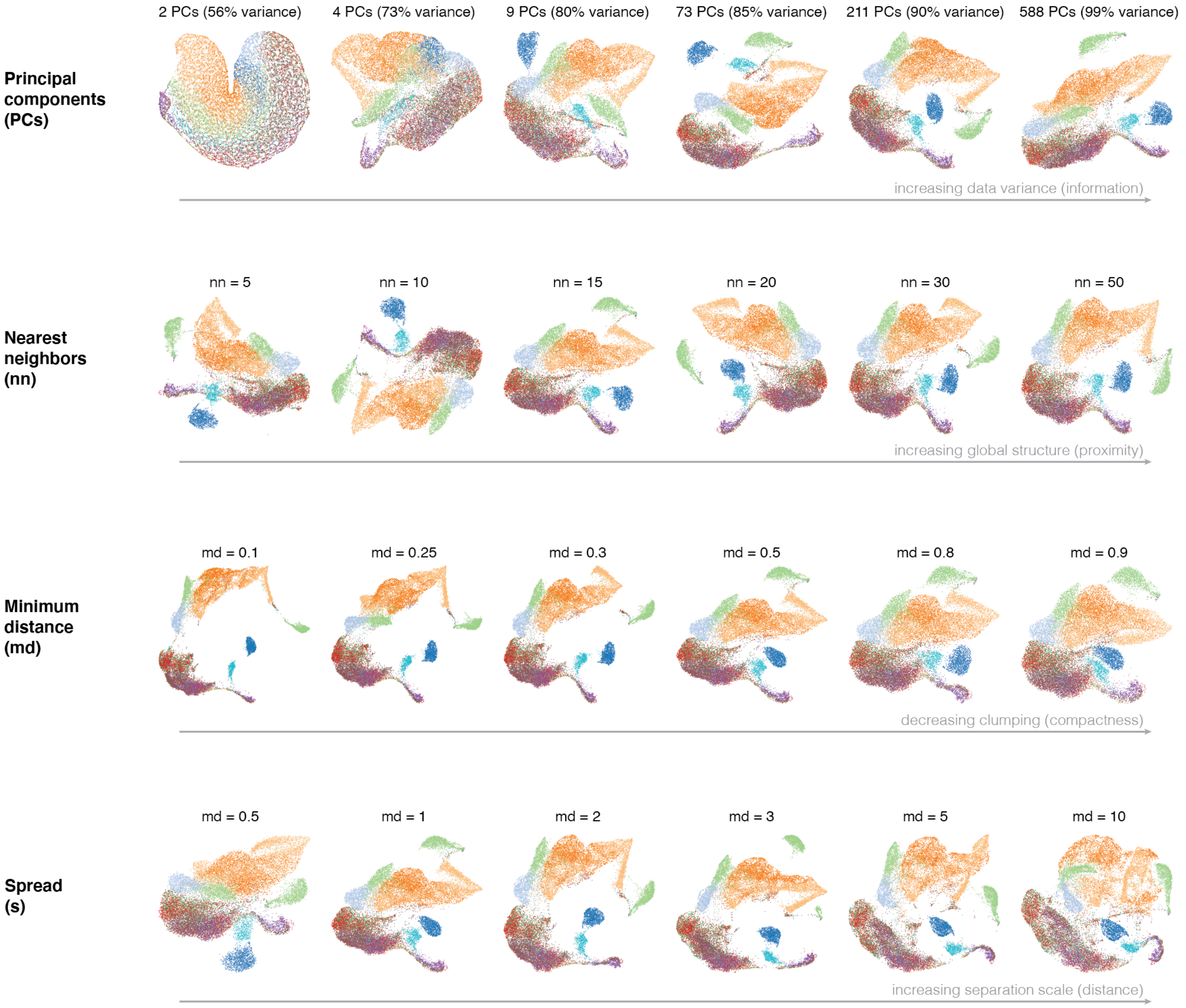


**Supplementary Figure 17. Hyperparameter effects on our data clustering and analysis.** Color corresponds to sample origin as defined in Figure 5 in the main text. Increasing the number of principal components retained more information and improved cell cluster separability. Increasing nearest neighbors examined more interconnectivity and preserved global structuring over local structures. Increasing minimum distance raised the lower bound of embedding space distance among points and decreased cell spectra crowding. Increasing spread enlarges the distribution of distances and affects the perceived distance between spectra.

a

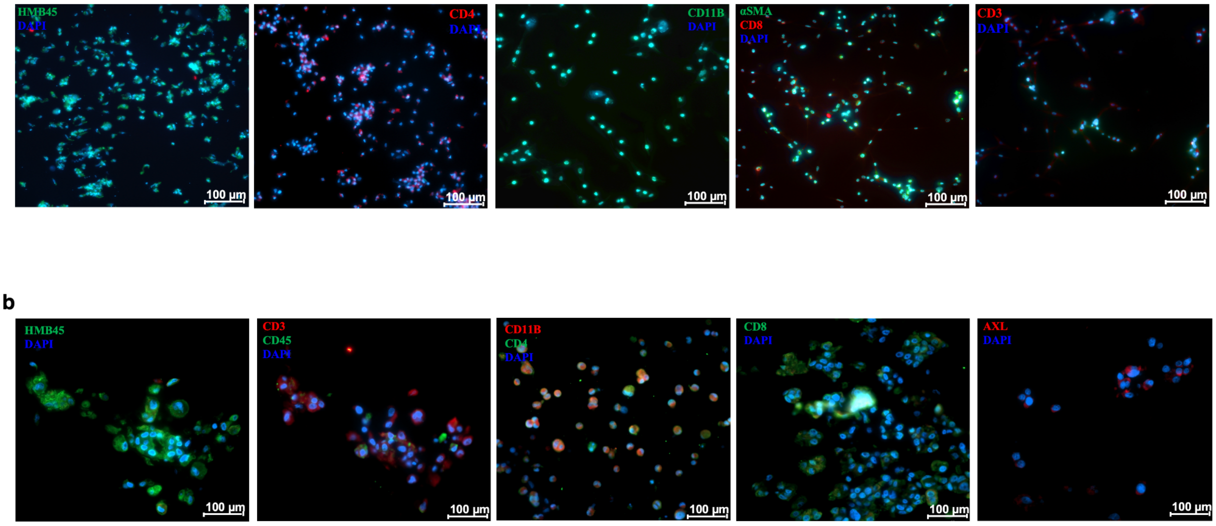


**Supplemental Figure 18. Immunofluorescence of patient samples.** Melanoma patient-derived samples retain immune cell composition and other cell types. A) PAT-52 patient-derived cells were allowed to grow for 7 days without passaging, and immunofluorescent tagging was applied with markers HMB-45, CD4, CD11B, α-SMA, CD8, and CD3. Scale bar, 100 μm. B) Images of PAT-73 patient-derived cells were allowed to grow for 7 days without passaging and immunofluorescent tagging was applied with markers HMB-45, CD3, CD45, CD11B, CD4, CD8, and AXL. Scale bar, 100 μm.

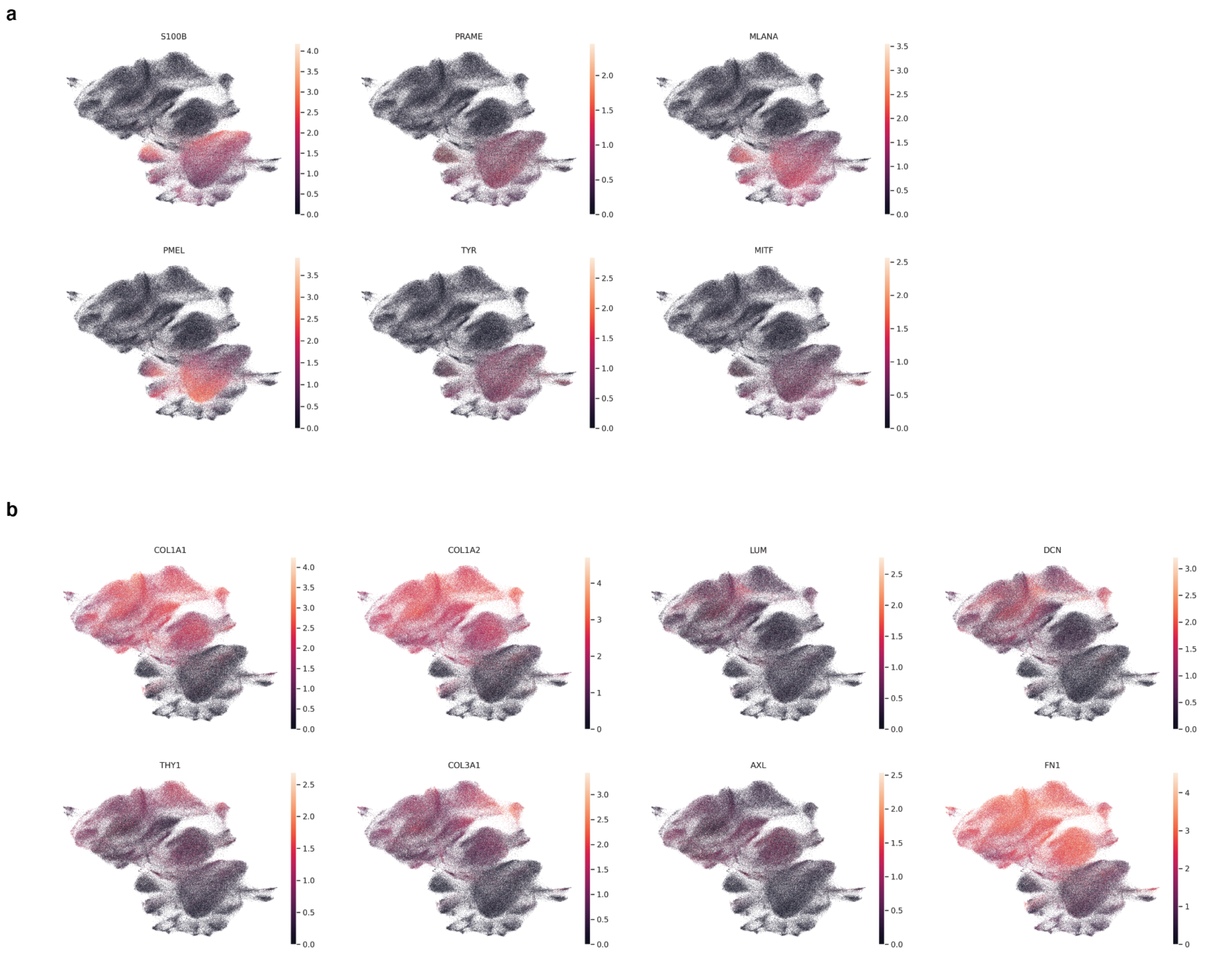


**Supplementary Figure 19. Feature plots of patient melanoma samples.** A) UMAP of our scRNA-seq dataset colored by marker genes S100B, PRAME, MLANA, PMEL, TYR, and MITF. The clusters with higher expressions of such genes are suggestive of tumor cells. B) UMAP of our scRNA-sequenced dataset colored by marker genes COL1A1, COL1A2, LUM, DCN, THY1, COL3A1, AXL, and PN1. The clusters with higher expressions of such genes are suggestive of fibroblast-like cells.

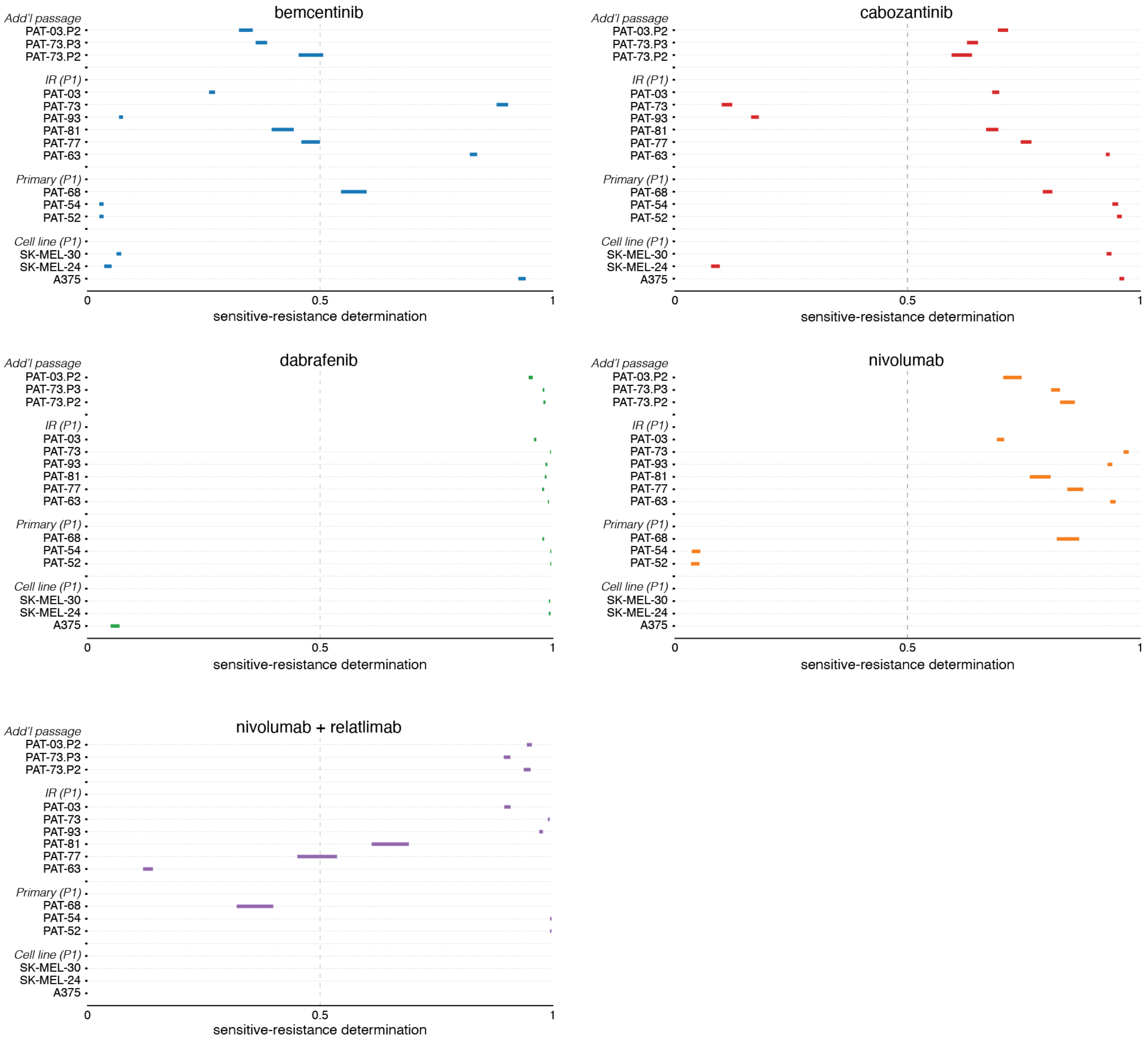


**Supplementary Figure 20. Percentile distribution of mini patient responses.** Response determination distribution for all mini patients sampled in each patient-derived cell line (seen and unseen by our ML model) across targeted and immunotherapy regimes. Bars represents the 5th to 95th percentile of the mini patient response.

### Supplementary Tables

| Mouse cell line viability | | | | |
|---|---|---|---|---|
| Cell line | Bem | Cabo | Dabra | Nivo |
| YUMM1.7 | 68.3 | 74.4 | 77.0 | 71.4 |
| YUMMER1.7 | 57.1 | 82.8 | 89.2 | 81.1 |

| Human cell line viability | | | | |
|---|---|---|---|---|
| Cell line | Bem | Cabo | Dabra | Nivo |
| A375 | 86.1 | 76.1 | 66.1 | 82.7 |
| SK-MEL-24 | 61.3 | 64.7 | 84.6 | 78.6 |
| SK-MEL-30 | 70.5 | 79.8 | 78.6 | 80.1 |

| Patient derived tumor line cell viability | | | | | | |
|---|---|---|---|---|---|---|
| Sample ID | Bem | Cabo | Dabra | Nivo | Nivo + Rela | Clinical |
| PAT-52 | 62.9 | 82.6 | 91.6 | 40.5 | 80.4 | - |
| PAT-54 | 70.4 | 72.4 | 89.6 | 45.6 | 85.4 | - |
| PAT-63 | 74.1 | 80.4 | 82.4 | 78.6 | 24.6 | NR (Nivo); R (Nivo + Rela) |
| PAT-73 | 92.0 | 71.2 | 92.1 | 86.7 | 90.6 | NR (Nivo) |
| PAT-93 | 52.4 | 69.4 | 82.4 | 66.1 | 90.4 | NR (Dabra); NR (Nivo) |
| PAT-68 | 83.4 | 85.2 | 74.8 | 86.4 | 27.4 | NR (Nivo) |
| PAT-03 | 62.7 | 65.4 | 81.9 | 66.1 | - | - |
| PAT-81 | 90.1 | 78.4 | 81.9 | 91.7 | 92.7 | NR (Dabra); NR (Nivo) |
| PAT-77 | 86.4 | 76.4 | 74.0 | 82.9 | 86.6 | NR (Nivo); NR (Nivo + Rela) |
| PAT-03.P2 | 72.7 | 73.4 | 80.4 | 70.1 | - | - |
| PAT-73.P2 | 89.0 | 61.2 | 81.1 | 85.7 | 89.0 | - |
| PAT-73.P3 | 71.0 | 77.2 | 89.1 | 90.7 | 91.1 | - |

**Supplementary Table 1. Cell viability data of mice and human melanoma samples.** In vitro cell viability assessments for the first passages were determined after 24 hour incubation of inhibitors with concentrations described in Table 3 to observe cell response for mice samples and human samples. Additionally, cell viability was determined for later passages on select patients of interest (PAT-73, PAT-03). Viability values correspond to relative percentage to untreated control samples. For the patient resistance determination model, resistant or sensitive labels were assigned based on cell viability inference or clinical outcomes in select cases. No clinical outcome labels (R for responder and NR for nonresponder) conflicted with cell viability observations.

| Inhibitor concentrations (μM) | | | | | |
|---|---|---|---|---|---|
| Sample | Bem | Cabo | Dabra | Nivo | Nivo + Rela |
| YUMM1.7 | 1 | 7 | 1 | 5 | - |
| YUMMER1.7 | 1 | 8.5 | 3.5 | 8 | - |
| A375 | 8 | 7.5 | 5 | 5 | - |
| SK-MEL-24 | 8 | 9.5 | 10 | 10 | - |
| SK-MEL-30 | 8.5 | 8.5 | 1 | 8 | - |
| Patient derived tumor lines | 8 | 9.5 | 5 | 6 | 6 + 7 |

**Supplementary Table 2.** Inhibitor concentrations used in each of our datasets.

| Cell response cell count | | | | | |
|---|---|---|---|---|---|
| Cell line | Control | Bem | Cabo | Dabra | Nivo |
| YUMM1.7 | 886 | 1025 | 1146 | 1081 | 1004 |
| YUMMER1.7 | 1078 | 961 | 958 | 838 | 960 |
| A375 | 1291 | 3103 | 2884 | 1267 | 1303 |
| SK-MEL-24 | 1299 | 1473 | 1423 | 1422 | 1430 |
| SK-MEL-30 | 3327 | 1506 | 1366 | 1680 | 1380 |
| PAT-52 | 1584 | 430 | 336 | 505 | 455 |
| PAT-73 | 2131 | 406 | 566 | 761 | 593 |

| Patient control cell count | |
|---|---|
| Sample ID | Count |
| PAT-52 | 1581 |
| PAT-54 | 1609 |
| PAT-63 | 557 |
| PAT-68 | 284 |
| PAT-73 | 2131 |
| PAT-77 | 96 |
| PAT-81 | 199 |
| PAT-93 | 657 |
| PAT-03 | 1512 |
| PAT-03.P2 | 1591 |
| PAT-73.P2 | 985 |
| PAT-73.P3 | 1388 |

| Patient control cell count | |
|---|---|
| Sample ID | Count |
| PAT-52.P2 | 1391 |
| PAT-54.P2 | 1386 |
| PAT-68.P2 | 1381 |
| PAT-63.P2 | 1329 |

**Supplementary Table 3.** Spectra count acquired for each sample utilized in our response or determination dataset. The first table represents spectra counts utilized in observing cell response to inhibitor presence. The second table represents spectra counts utilized in our patient prediction model analysis. The third table represents spectra counts from additional passages of select patients used in understanding passage shift.

**Supplementary Table 4.** Raman band assignments associated with tumor immune microenvironment cells.

| Raman band ($cm^{-1}$) | Band assignment | Biological group | Reference |
|---|---|---|---|
| 644-645 | C-C twisting in Tyrosine | Proteins | [1–4] |
| 654 | Tyrosine, G | Proteins, Nucleic acid | [5] |
| 667 | T, G | Nucleic acids | [2,6] |
| 708 | C | Nucleic acids | [7,8] |
| 731 | A (ring breathing) | Nucleic acids | [2] |
| 745-750 | Ring breathing vibrations of nucleic acids<br>Tryptophan | Nucleic acids<br>Proteins | [9] |
| 779-784 | C, U (ring breathing), PO2- backbone stretch | Nucleic acids | [2,10] |
| 805 | PO2- symmetric stretch | Nucleic acids | [11] |
| 826 | Tyrosine (ring breathing) | Proteins | [12] |
| 829 | Ring breathing | Proteins | [13,14] |
| 833-836 | PO2- symmetric stretch | Nucleic acids | [12] |
| 841-843 | Polysaccharide, Glucose | Carbohydrates | [6] |
| 845-851 | C-C stretch in Tyrosine and Valine | Proteins | [4,13,15] |
| 876 | Tryptophan | Proteins | [14,16] |
| 883 | CH2 rocking | Proteins | [6] |
| 888 | Tyrosine | Proteins | [17] |
| 904 | Tyrosine<br>Glycogen | Proteins<br>Carbohydrates | [18,19] |
| 912 | Glucose | Carbohydrates | [6,20] |
| 917 | C-C stretch | Proteins | [20] |
| 924 | Proline ring | Protein | [11] |
| 927-940 | C-C skeletal stretch | Proteins | [6,12,21] |
| 955 | C-C stretch | Proteins | [3] |
| 958-959 | CH3 deformations | Lipids, Proteins | [22] |
| 968 | Phosphatidylcholine | Lipids | [12] |
| 998-1003 | C-C aromatic symmetric ring breathing of phenylalanine | Proteins | [23–25] |

| | | | |
|---|---|---|---|
| 1021 | Glycogen | Carbohydrates | [6] |
| 1030-1032 | C-N in-plane bending of phenylalanine | Proteins | [12] |
| 1080-1082 | C-C<br>PO2 symmetric stretch | Lipids<br>Nucleic acids | [26,27] |
| 1126-1129 | C–C stretch<br>C–N stretch | Lipids<br>Proteins | [1,25] |
| 1156-1158 | C-C stretch, C-N stretch | Beta-carotene<br>Proteins | [4,27–29] |
| 1173-1174 | Tyrosine, Phenylalanine | Proteins | [2] |
| 1205-1207 | C-C6H5 stretch in phenylalanine and tryptophan<br>Tyrosine | Proteins | [4,12,30] |
| 1220 | Amide III | Proteins | [31] |
| 1225 | Amide III | Proteins | [6] |
| 1241 | T, Amide III | Nucleic acids<br>Proteins | [2] |
| 1250-1252 | Amide III | Proteins | [12] |
| 1270 | C-H bending | Lipids | [32] |
| 1274 | Amide III | Proteins | [33] |
| 1303-1306 | A (ring mode),<br>C-H deformation<br>CH2 twist | Nucleic acids<br>Proteins<br>Lipids | [2,12,34,35] |
| 1308-1310 | CH2 deformation, CH3CH2 twisting | Lipids | [9,30] |
| 1312 | CH3CH2 twisting | Proteins, Lipids | [31] |
| 1336 | CH3, CH2 wagging, C-H deformation, C-C stretch | Proteins, Lipids,<br>Carbohydrates | [15,36] |
| 1337 | A, G (ring mode) | Nucleic acids | [29] |
| 1352 | G<br>Tryptophan | Nucleic acids<br>Proteins | [37] |
| 1361 | Tryptophan | Proteins | [20] |
| 1382 | CH3 symmetric deformation<br>Ring breathing modes of the nucleic acids | Lipids<br>Nucleic acids | [38] |
| 1385 | C-C linear stretch<br>CH3 symmetric deformation | Melanin<br>Lipids | [39–42] |
| 1400 | U, A | Nucleic acids | [8] |
| 1410-1413 | CH deformation | Proteins<br>Lipids | [43] |

| | | | |
|---|---|---|---|
| 1419-1420 | CH2 scissoring<br>A, G | Lipids<br>Nucleic acids | [12] |
| 1433-1440 | CH2 deformation, C=C stretch unsaturated fatty acids | Lipids | [9,10] |
| 1445-1447 | CH2 bend<br>CH2 deformation | Proteins<br>Lipids | [2,10,25] |
| 1457-1460 | CH2 deformation, CH deformation | Proteins | [2,12] |
| 1470 | G, A | Nucleic acid | [44] |
| 1482 | G, A (ring mode) | Nucleic acids | [12] |
| 1497 | C-NH2 scissoring | Nucleic acids | [45] |
| 1518 | C=C stretch | Carotenoid | [6] |
| 1526-1527 | C=C | Carotenoids | [2] |
| 1553 | Tryptophan | Proteins | [12] |
| 1562 | C-N and N-H vibrations<br>Tryptophan<br>A, G ring breathing | Proteins<br>Nucleic acids | [38] |
| 1575 | C-C ring stretch | Nucleic acids | [12,15] |
| 1580 | Aromatic ring in-plane stretch | Melanin | [39] |
| 1582-1588 | A,G<br>C=C olefinic stretch<br>Tyrosine | Nucleic acids<br>Proteins | [6,9,13,27,39–41,46] |
| 1591-1593 | C=C in-plane bending mode in Phenylalanine | Proteins | [42] |
| 1606 | C=C stretch in Phenylalanine | Proteins | [47] |
| 1610 | aromatic ring stretch in Tyrosine | Proteins | [4] |
| 1624 | C=C bend in tryptophan | Proteins | [45] |
| 1636 | Amide I | Proteins | [6] |
| 1652-1658 | C=C stretch, Amide I | Lipids, Proteins | [9,29,42] |
| 1660-1666 | Amide I | Proteins | [12,30] |
| 1672 | Amide I | Proteins | [15,33] |
| 1681-1684 | U (C=O stretch) | Nucleic acids | [12] |
| 1692 | Amide I | Proteins | [7] |
| 1735-1775 | C=O stretch | Lipids | [27,42,48] |
| 1791-1797 | | Unassigned | [49] |

24. Ribeiro, A. R. B. *et al.* Application of Raman spectroscopy for characterization of the functional polarization of macrophages into M1 and M2 cells. *Spectrochim Acta A Mol Biomol Spectrosc* **265**,

120328 (2022).
